# Assessment of computational methods for the analysis of single-cell ATAC-seq data

**DOI:** 10.1101/739011

**Authors:** Huidong Chen, Caleb Lareau, Tommaso Andreani, Michael E. Vinyard, Sara P. Garcia, Kendell Clement, Miguel A Andrade-Navarro, Jason D. Buenrostro, Luca Pinello

**Affiliations:** Molecular Pathology Unit, Massachusetts General Hospital Research Institute, Charlestown, MA 02129, USA; Center for Cancer Research, Massachusetts General Hospital, Charlestown, MA 02129, USA; Department of Pathology, Harvard Medical School, Boston, MA 02115, USA; Broad Institute of Harvard and MIT, Cambridge, MA 02142, USA; Department of Stem Cell and Regenerative Biology, Harvard University, Cambridge, MA 02138, USA; Faculty of Biology, Computational Biology and Data Mining Lab. Johannes Gutenberg University of Mainz, 55128 Mainz, Germany; Department of Chemistry and Chemical Biology, Harvard University, Cambridge, MA 02142, USA

**Author notes:** These authors contributed equally (Caleb Lareau, Tommaso Andreani, Michael E. Vinyard). Correspondence should be addressed to L.P.

**Keywords:** scATAC-seq, feature matrix, benchmarking, regulatory genomics, clustering, visualization, featurization, dimensionality reduction

## Abstract

**Background:** Recent innovations in single-cell Assay for Transposase Accessible Chromatin using sequencing (scATAC-seq) enable profiling of the epigenetic landscape of thousands of individual cells. scATAC-seq data analysis presents unique methodological challenges. scATAC-seq experiments sample DNA, which, due to low copy numbers (diploid in humans) lead to inherent data sparsity (1-10% of peaks detected per cell) compared to transcriptomic (scRNA-seq) data (20-50% of expressed genes detected per cell). Such challenges in data generation emphasize the need for informative features to assess cell heterogeneity at the chromatin level.

**Results:** We present a benchmarking framework that was applied to 10 computational methods for scATAC-seq on 13 synthetic and real datasets from different assays, profiling cell types from diverse tissues and organisms. Methods for processing and featurizing scATAC-seq data were evaluated by their ability to discriminate cell types when combined with common unsupervised clustering approaches. We rank evaluated methods and discuss computational challenges associated with scATAC-seq analysis including inherently sparse data, determination of features, peak calling, the effects of sequencing coverage and noise, and clustering performance. Running times and memory requirements are also discussed.

**Conclusions:** This reference summary of scATAC-seq methods offers recommendations for best practices with consideration for both the non-expert user and the methods developer. Despite variation across methods and datasets, SnapATAC, *Cusanovich2018*, and cisTopic outperform other methods in separating cell populations of different coverages and noise levels in both synthetic and real datasets. Notably, SnapATAC was the only method able to analyze a large dataset (> 80,000 cells).

## Background

Individual cell types within heterogenous tissues coordinate to perform complex biological functions, many of which are not fully understood. Recent technological advances in single-cell methodologies have resulted in an increased capacity to study cell-to-cell heterogeneity and the underlying molecular regulatory programs that drive such variation.

To date, most single-cell profiling efforts have been performed via quantification of RNA by sequencing (scRNA-seq). While this provides snapshots of inter- and intra-cellular variability in gene expression, investigation of the *epigenomic* landscape in single cells holds great promise for uncovering an important component of the regulatory logic of gene expression programs. Enabled by advances in array-based technologies, droplet microfluidics and combinatorial indexing through split-pooling[1] (**Fig. 1a**), single-cell Assay for Transposase Accessible Chromatin using sequencing (scATAC-seq) has recently overcome previous limitations of technology and scale to generate chromatin accessibility data for thousands of single cells in a relatively easy and cost-effective manner.

**Figure 1.**
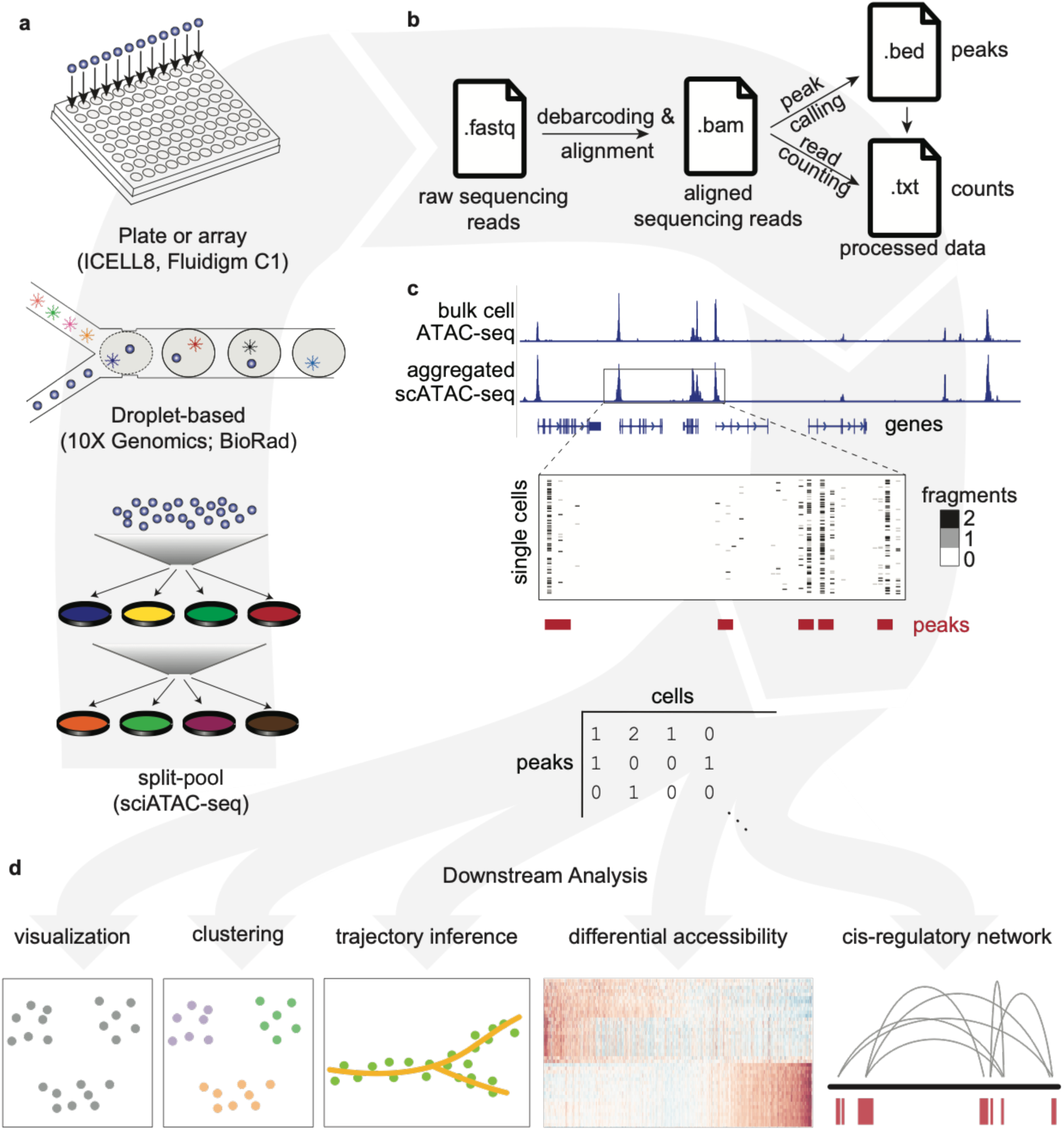
Schematic overview of single cell ATAC-seq assays and analysis steps. (a) Single cell ATAC libraries are created from single cells that have been exposed to the Tn5 transposase using one of three protocols: 1) Single cells are individually barcoded by a split-and-pool approach where unique barcodes added at each step can be used to identify reads originating from each cell 2) microfluidic droplet-based technologies provided by 10x Genomics and BioRad are used to extract and label DNA from each cell or 3) each single cell is deposited into a multi-well plate or array from ICELL8 or Fluidigm C1 for library preparation. (b) After sequencing, the raw reads obtained in .fastq format for each single cell are mapped to a reference genome, producing aligned reads in .bam format. Finally, peak calling and read counting return the genomic position and the read count files in. bed and .txt format, respectively. Data in these file formats is then used for downstream analysis. (c) ATAC-seq peaks in bulk samples can generally be recapitulated in aggregated single cell samples, but not every single cell has a fragment at every peak. A feature matrix can be constructed from single cells (e.g., by counting the number of reads at each peak for every cell). (d) Following construction of the feature matrix, common downstream analyses including visualization, clustering, trajectory inference, determination of differential accessibility, and the prediction of cis-regulatory networks can be performed using the methods benchmarked in this manuscript.

However, the analysis of scATAC-seq data presents methodological challenges distinct from those of single-cell transcriptomic (scRNA-seq) data. The primary difficulty arises from a difference in the number of RNA vs DNA molecules available for profiling in single cells. While for an expressed gene several RNA molecules are present in a single cell, scATAC-seq assays profile DNA, a molecule which is present in only few copies per cell (two in a diploid organism). The low copy number results in an inherent per-cell data sparsity, where only 1-10% of expected accessible peaks are detected in single cells from scATAC-seq data, compared to 20-50% of expressed genes detected in single cells from scRNA-seq data. This emphasizes the need to recover informative features from sparse data to assess variability between cells in scATAC-seq analyses. Further, determination of which features best define cell state is currently unclear.

The difference in readout (gene expression versus chromatin accessibility) has also motivated a variety of approaches to selecting informative features in scATAC-seq methods. While most processing pipelines share common upstream processing steps (i.e. alignment, peak calling, and counting; **Fig. 1b**), existing computational approaches differ in the way they obtain a feature matrix for downstream analyses. For example, some methods select features based on the sequence content of accessible regions (e.g. *k*-mer frequencies[2, 3] or transcription factor (TF) motifs [3]), whereas other methods select features based on the genomic coordinates of the accessible regions (e.g. extended promoter regions to determine chromatin activity surrounding genes [2, 4]). Finally, the potential feature set in scATAC-seq, which includes genome-wide regions of accessible chromatin (**Fig. 1c**), is typically 10-20x the size of the feature set in scRNA-seq experiments (which is defined and limited by the number of genes expressed). This larger feature set could be valuable in distinguishing a wider variety of cell populations and inferring the dynamics underlying cell organization into complex tissues[5].

However, the novelty and assay-specific challenges associated with these large-scale scATAC-seq datasets and the lack of analysis guidelines have resulted in diverging computational strategies to aggregate data across such an immense feature space with no clear indication as to which strategy or strategies are most advantageous.

Here, we provide the first benchmark assessment of computational methods for the analysis of scATAC-seq data. We discuss the impact of feature matrix construction strategies (e.g. sequence content-based vs. genomic coordinates) on common downstream analysis, with a focus on clustering and visualization. This comprehensive survey of current available methods provides user-specific recommendations for best practices that aim to maximize inference-capability for current and future scATAC-seq workflows. Importantly, we provide more than 100 well-documented Jupyter Notebooks (https://github.com/pinellolab/scATAC-benchmarking/) to easily reproduce our analyses. We anticipate that this will be a valuable resource for future scATAC-seq benchmark studies.

## Results

### Benchmark Framework

For this benchmarking study we created an unbiased framework to qualitatively and quantitatively survey the ability of available scATAC-seq methods to featurize chromatin accessibility data. Evaluated using this framework were several datasets of divergent size and profiling technologies. Using widely accepted quantitative metrics, we explored how differences in feature matrix construction influence outcomes in exploratory visualization and clustering, two common downstream analyses. The general overview of our framework is presented in **Fig. 2**.

**Figure 2.**
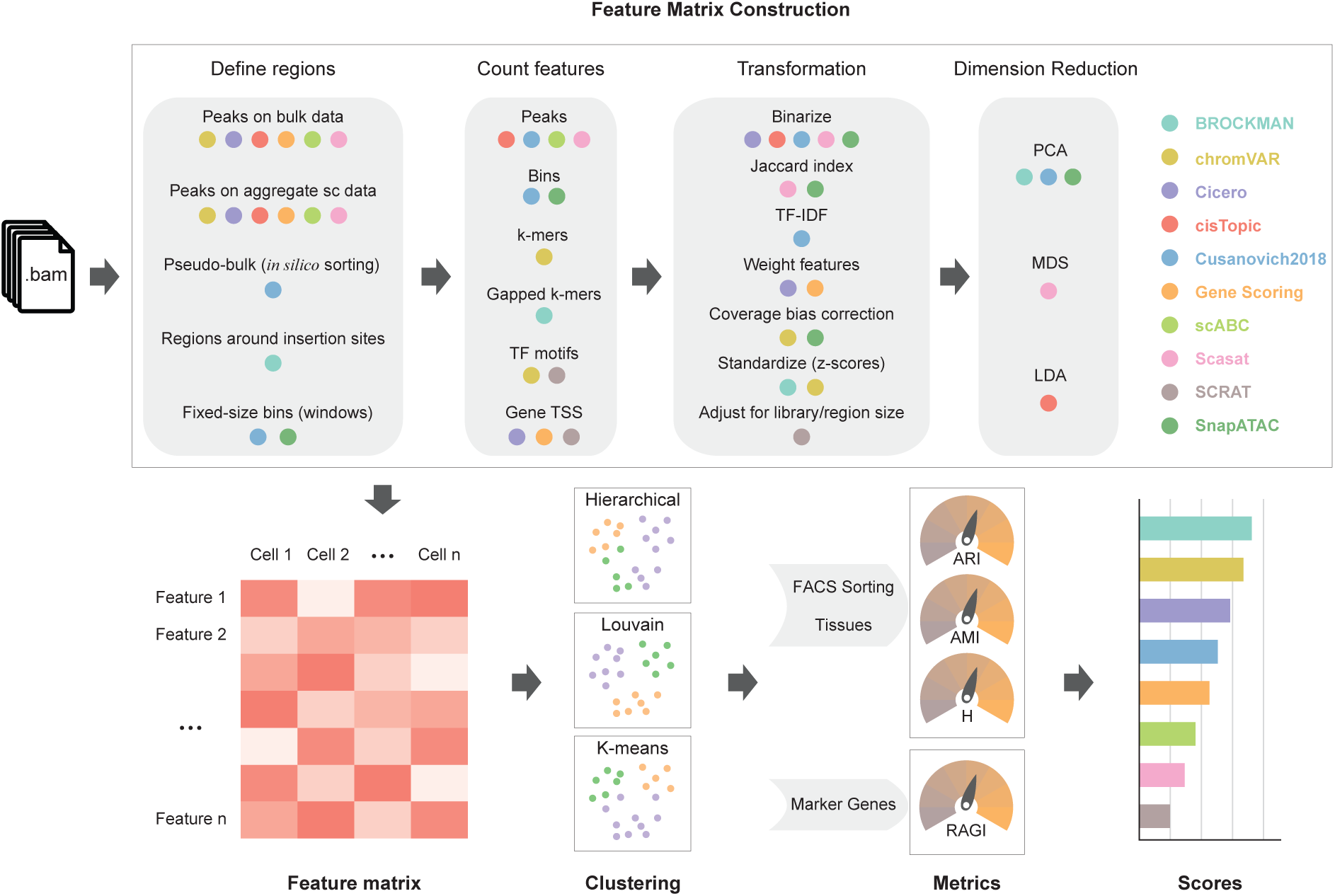
Benchmarking workflow. Starting from aligned read files in .bam format, feature matrices were constructed using each method. The feature matrix construction techniques used by each method were grouped into four broad categories: *Define regions*, *Count features*, *Transformation* and *Dimension Reduction*. A colored dot under a technique indicates that the method (signified by the respective color in the legend on the right) uses that technique. For each method, feature matrix files (defined as columns as cells and rows as features) are calculated and used to perform hierarchical, Louvain and k-means clustering analysis. For datasets with a ground truth such as FACS-sorting labels or known tissues, clustering evaluation was performed according to the Adjusted Random Index (ARI), Adjusted Mutual Information (AMI) and homogeneity (H) scores. For datasets without ground truth, the clustering solutions were evaluated according to a Residual Average Gini Index (RAGI), a metric that compares cluster separation based on known marker genes against housekeeping genes. Lastly, a final score is assigned to each method.

For this study we collected public data from three published studies (aligned files in BAM format) and generated ten simulated datasets with various coverages and noise levels (see **Methods**). To calculate feature matrices for downstream analysis, for each method we followed the guidelines provided in the documentation in the original study or as suggested by the respective authors. After feature matrix construction, we used three commonly used clustering approaches (K-means, Louvain and Hierarchical Clustering)[6] and UMAP[7] projection to find putative subpopulations and visualize cell-to-cell similarities for each method. Next, the quality of the clustering solutions was evaluated by adjusted random index (ARI), adjusted mutual information (AMI) and homogeneity (H) when FACS-sorting labels or tissues were available (gold standard); or by a proposed Gini-index-based metric called Residual Average Gini Index (RAGI) when only known marker genes were available (silver standard). Finally, based on these metrics, the methods were ranked by the quality of their clustering solutions across datasets.

### Methods overview and featurization of chromatin accessibility data

Several computational methods have been developed to address the inherent sparsity and high dimensionality of single cell ATAC-seq data, including BROCKMAN[3], chromVAR[2], Cicero[8], cisTopic[9], *Cusanovich2018*[1, 10, 11], Gene Scoring[12], scABC[13], Scasat[14], SCRAT[4], and SnapATAC[15]. Based on the proposed workflow of each method, we were able to compute different feature matrices defined as a features-by-cells matrix (e.g. read counts for each cell (columns) in a given open chromatin peak *feature* (rows)) that could then be readily used for downstream analyses such as clustering. Starting from single cell BAM files, the feature matrix construction can be roughly summarized into four different common modules: *define regions*, *count features*, *transformation*, *and dimensionality reduction* as illustrated in **Fig. 2**. Not every method uses all steps, therefore we provide below, a short summary of the strategies adopted by each method and a *per module* discussion to highlight key similarities and differences (for a more detailed description of each strategy see **Methods**).

Briefly, BROCKMAN[3] represents genomic sequences by gapped k-mers (short DNA sequences of length k) within transposon integration sites and infers the variation in k-mer occupancy using principal component analysis (PCA). chromVAR[2] estimates the dispersion of chromatin accessibility within peaks sharing the same feature, e.g. motifs or k-mers. Cicero[8] calculates a gene activity score based on accessibility at a promoter region and the regulatory potential of peaks nearby. cisTopic[9] applies Latent Dirichlet Allocation (LDA) (a Bayesian topic modeling approach commonly used in natural language processing) to identify cell states from topic-cell distribution and explore cis-regulatory regions from region-topic distribution. Previous approaches that utilize latent semantic indexing (LSI) (termed here as *Cusanovich2018*)[1, 10, 11] first partition the genome into windows, normalize reads within windows using the term frequency-inverse document frequency transformation (TF-IDF), reduce dimensionality using singular value decomposition (SVD), and perform a first-round of clustering (referred to as ‘in silico cell sorting’) to generate clades and call peaks within them. Finally, the clusters are refined with a second-round of clustering after TF-IDF and SVD based on read counts in peaks. The Gene Scoring method[12] assigns each gene an accessibility score by summarizing peaks near its transcription start site (TSS) and weighting them by an exponential decay function based on their distances to the TSS. scABC[13] first calculates a global weight for each cell by taking into account the number of distinct reads in the regions flanking peaks (to estimate the expected background). Based on these weights, it then uses weighted k-medoids to cluster cells based on the reads in peaks. Scasat[14] binarizes peak accessibility and uses multidimensional scaling (MDS) based on the Jaccard distance to reduce dimensionality before clustering. SCRAT[4] summarizes read counts on different regulatory features (e.g. transcription factor binding motifs, gene TSS regions). SnapATAC[15] segments the genome into uniformly-sized bins and adjusts for differences in library size between cells using a regression-based normalization method; finally PCA is performed to select the most significant components for clustering analysis.

#### Define Regions

An essential aspect of feature matrix construction is the selection of a set of regions to describe the data (e.g. putative regulatory elements such as peaks, promoters etc.). Most methods described above, including chromVAR, Cicero, cisTopic, Gene Scoring, scABC, and Scasat, define regions based on peak calling from either a reference bulk ATAC-seq profile or an aggregated single cell ATAC-seq profile. *Cusanovich2018*, as briefly mentioned above, instead of aggregating single cell to call peaks, first creates pseudo-bulk clades by performing hierarchical clustering on the TF-IDF and SVD transformed matrix using the top frequently accessible windows. Then peaks are called by aggregating cells within each pseudo-bulk clade. In addition to relying on peaks, some methods have proposed different strategies. BROCKMAN uses the union of regions around transposon integration sites. *Cusanovich2018* (before *in silico* sorting) and SnapATAC segment the genomes into fixed-size bins (windows) and count features within each bin.

#### Count Features

Once feature regions are defined, raw features within these regions are counted. Note that some methods (e.g. chromVAR) may support the counting of multiple features. For cisTopic, *Cusanovich2018*, scABC, and Scasat, reads overlapping peaks are counted. For *Cusanovich2018* (before the *in silico* sorting step) and SnapATAC, reads overlapping bins are counted. k-mers are counted under peaks for chromVAR while gapped k-mers are counted for BROCKMAN around transposase cut sites. Similarly, transcription factor motifs (e.g. from the JASPAR database[16]) can be used as features by counting reads overlapping their binding sites in peaks (chromVAR) or genome-wide (SCRAT). If predefined genomic annotations such as coding genes are given, Gene Scoring, Cicero, and SCRAT use gene TSSs as anchor points to calculate gene enrichment scores based on reads nearby or just within peaks nearby.

#### Transformation

After building the initial raw feature matrix using the counting step, different transformation methods can be performed. Binarization of read counts is used by five out of the ten evaluated methods: Cicero, cisTopic, *Cusanovich2018*, Scasat, and SnapATAC. (**Fig. 2**). This step is based on the assumption that each site is present at most twice (for diploid genomes) and that the count matrix is inherently sparse. Binarization is advantageous in alleviating challenges arising from sequencing depth or PCR amplification artifacts. SnapATAC and Scasat convert the binary count matrix into a cell-pairwise Jaccard index similarity matrix. *Cusanovich2018* normalizes the binary count matrix using the TF-IDF transformation. Cicero weights feature sites by their co-accessibility, while Gene Scoring weights sites by a decaying function based on its distance to a gene TSS. Both chromVAR and SnapATAC perform a read coverage bias correction to account for the influence of sample depth. scABC also implements a similar step but calculates a weight for each cell; even if these weights are not used to transform the matrix, they are used later in the clustering procedure. SCRAT adjusts for both library size and region length. chromVAR creates ‘background’ peaks consisting of an equal number of peaks matched for both average accessibility and GC content to calculate bias-corrected deviation. Both BROCKMAN and chromVAR compute z-scores to measure the gain or loss of chromatin accessibility across cells.

#### Dimensionality Reduction

In the final step before downstream analysis, several methods apply different dimensionality reduction techniques to project the cells into a space of fewer dimensions. This step can refine the feature space mitigating redundant features and potential artifacts, and potentially reducing the computation time of downstream analysis (**Fig. 2**). PCA is the most commonly used method (used by BROCKMAN, SnapATAC, and *Cusanovich2018*). cisTopic uses latent Dirichlet allocation (LDA) to generate two distributions including topic-cell distribution and region-topic distribution. Choosing the top topics based on the topic-cell distribution reduces the dimensionality. Scasat uses multidimensional scaling (MDS). When reviewing the different methods to include in our benchmark, we noticed that not all methods perform a dimensionality reduction step, which could skew the relative performance across methods. Therefore, for chromVAR, Cicero (gene activity score), Gene Scoring, scABC, and SCRAT, we considered in addition to the original feature matrix, also a new feature matrix after PCA transformation, since this is simple and commonly used technique for dimensionality reduction.

To better evaluate the effects of different modules including *define regions*, *count features*, *transformation*, and *dimensionality reduction*, we also considered a simple control method, referred to as Control-Naïve, by combining the most common and simple steps for building a feature matrix, i.e. counting reads within peaks to obtain a peaks-by-cells raw count matrix and then performing PCA on it (the number of top principal components was determined based on the elbow plot for all the methods). Since the feature matrix of scABC is also a peaks-by-cells raw count matrix, this matrix after PCA will correspond to the one obtained by the Control-Naïve method (to avoid redundancies, in our assessment we refer to this matrix as Control-Naïve).

We also noticed that some methods might slightly diverge from the proposed four modules common framework. For example, Cicero calculates gene activity scores by first performing two transformations (binarize and weight features) and then performing the counting step around the annotated TSS. We believe the proposed modularization of the of the feature matrix construction can still serve as a useful framework to represent the core components of the different methods and provides an intuitive and informative summary of the diverse scATAC-seq methodologies.

Once dimensionality reduction is completed, the transformed feature matrix can be used for unbiased clustering, visualization, or other downstream analyses. Here we have used the final feature matrices generated by each scATAC-seq analysis method, and evaluated their performance in uncovering different populations by unsupervised clustering.

### Clustering approaches and metrics used for performance evaluation

This study employed three diverse types of commonly used unsupervised clustering methods for single cell analysis [6]: K-means clustering, Hierarchical Clustering, and the Louvain community detection algorithm (see **Methods**).

Clustering results were evaluated by three commonly used metrics: adjusted random index (ARI), adjusted mutual information (AMI) and homogeneity when a gold standard solution was available (known labels for the simulation data and FACS-sorted cell populations or known tissues for the real datasets). We propose a Gini-index-based metric called Residual Average Gini Index (RAGI), which was used to evaluate the clustering results when no ground truth was available and only a few marker genes were known by which populations could be discriminated (see **Methods**). For each metric, we defined the *clustering score* as the highest score amongst the three clustering methods, i.e. the score which corresponded to the clustering solution that maximized the metric.

This framework allowed for benchmarking the ability of each strategy to featurize chromatin accessibility data and its impact on important downstream analyses such as clustering and visualization. The following sections present the results of this evaluation for all above-described synthetic and real scATAC-seq datasets.

### Clustering performance on simulated datasets

We simulated 10 scATAC-seq datasets using available bulk ATAC-seq datasets with clear annotations from bone marrow and erythropoiesis[5, 17] using varying noise levels and read coverages. Briefly, to generate the peak by cell matrices, we defined a noise parameter (between 0 and 1) as the proportion of reads occurring in a random peak from one of the sorted populations. The remaining proportion of reads was distributed as a function of the bulk sample (see Methods). A feature matrix with a noise level of 0 preserved perfectly the underlying cell type specificity of the reads within peaks. Conversely, a feature matrix with a noise level of 1, contained no information to discriminate cell types based on the reads within peaks. In our study, we considered three noise levels: no noise (0), moderate noise (0.2) and high noise (0.4). To better and more fairly evaluate the contribution of the core steps of each method (i.e. *count features*, *transformation* and *dimensionality reduction*) regardless of the preprocessing steps usually excluded from these methods (reads filtering, alignment, peak calling, etc.), we compared the performance of each method using a set of predefined peak regions from bulk ATAC-seq datasets. We selected the top 80,000 peaks based on the number of cells in which peaks were observed (each peak that was present in at least one cell) for all methods and all synthetic datasets.

Using the bulk ATAC-seq bone marrow dataset, we simulated five additional datasets to explore the effect of coverage on clustering performance (5,000 fragments, 2,500 fragments, 1,000 fragments, 500 fragments, 250 fragments respectively per cell).

Each method was used to analyze all synthetic datasets as suggested in the method documentation (see **Sup Note 1** and **Sup Fig. 1**).

#### Simulated bone marrow datasets

We generated chromatin accessibility profiles (2,500 fragments per cell) based on six different FACS-sorted bulk cell populations: hematopoietic stem cells (HSCs), common myeloid progenitor cells (CMPs), erythroid cells (Ery), and other three lymphoid cell types: natural killer cells (NK), CD4 and CD8 T-cells (see **Fig. 3a**). We used ARI, AMI and homogeneity metrics to compare the clustering solutions with the known cell type labels (**Fig. 3b, Sup Fig. 2, Sup Table 1**). The top three methods based on these simulation settings were cisTopic, *Cusanovich2018*, and SnapATAC. They performed equally well with no noise and moderate noise (with clustering scores close to 1.0) (**Sup Fig. 2, Sup Table 2**). At a noise level of 0.4, the methods showed more separation in performance accordingly to the three metrics (**Fig. 3b, Sup Table 3**). SnapATAC, *Cusanovich2018*, and cisTopic clearly outperformed the Control-Naïve method with consistently higher clustering scores across all metrics. Scasat performed slightly better than the Control-Naïve method, and the remaining methods under-performed relative to the Control-Naïve method. For scABC (i.e. peaks-by-cells raw count matrix), Hierarchical Clustering performs much better than the other two clustering methods. chromVAR performance using k-mers as features was superior to the approach using motifs. Another k-mer-based method, BROCKMAN demonstrated similar performance to the k-mer-based chromVAR method. Motif-based SCRAT performed better than motif-based chromVAR. Both Cicero gene activity scores and Gene Scoring (which summarize the chromatin accessibility around coding annotations without a dimensionality reduction step) generally performed poorly. PCA boosted performance of scABC, Cicero, and Gene Scoring. This step improved clustering performance regardless of the clustering method (also we noted again that scABC after PCA is equivalent to the Control-Naïve method), especially for the Louvain approach. PCA also slightly boosted performance of the k-mer-based chromVAR but did not markedly improve the results of the motif-based chromVAR or SCRAT analyses.

**Figure 3.**
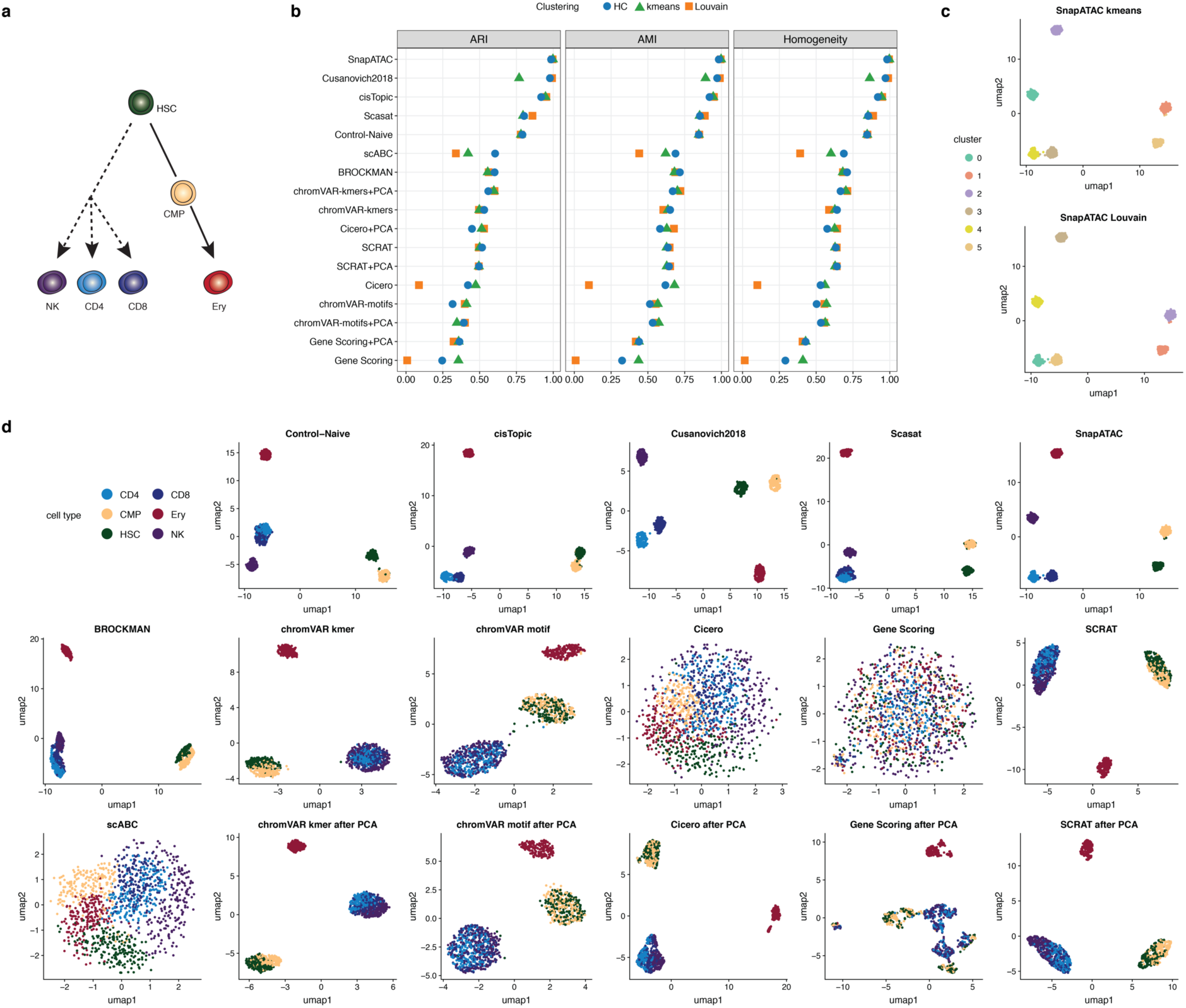
Benchmarking results in simulated bone marrow datasets at a noise level of 0.4 and a coverage of 2,500 fragments. **(a)** Cell types used to create the simulated dataset. **(b)** Dot plot of scores for each metric to quantitatively measure the clustering performance of each method, sorted by maximum ARI score. **(c)** The two top-scoring pairings of scATAC-seq analysis method and clustering technique. Cell cluster assignments from each method are shown using the colors in the legend on the left. **(d)** UMAP visualization of the feature matrix produced by each method for the simulated dataset. Individual cells are colored indicating the cell type labels shown in (a).

We next investigated qualitatively the obtained clustering solutions, using the respective feature matrices to project the cells onto a 2-D space using UMAP and colored them based on the obtained clustering solutions (**Sup Fig. 3**) or based on the true population labels used to generate the data (**Fig. 3d**). The top two clustering solutions based on the ARI (SnapATAC with k-means and SnapATAC with Louvain) are shown for ease of comparison (**Fig. 3c**).

*Cusanovich2018* and SnapATAC are the only two methods that clearly separated all six populations. cisTopic slightly mixed CD4 and CD8 T-cells. Scasat and the Control-Naïve method failed to separate CD4 and CD8 T-cell populations. BROCKMAN slightly mixed NK with CD4 and CD8 T-cells and could not further separate CD4 and CD8 T-cells. It also failed to clearly separate HSC and CMP. Both kmer-based and motif-based chromVAR as well as SCRAT could only separate the Ery population while failing to separate HSC and CMP as well as CD4, CD8 T-cells, and NK. The chromVAR k-mers-based method mixed HSC and CMP to a lesser extent compared to the motifs-based method. There was no clear separation of cells using scABC (the peaks-by-cells raw count matrix), Cicero, or Gene Scoring. We observed that PCA clearly improved the separation of cell populations for Cicero and Gene Scoring. It also slightly improved the separation of CD4, CD8 T-cells, and NK populations by k-mer-based chromVAR. No clear improvement was observed for the motif-based chromVAR, or SCRAT methods. We further observed that a lack of visual separation of cell types in the UMAP plots (scABC, Cicero, and Gene Scoring), corresponded with substantial variation between the performances of the three clustering methods, showing better performance in the k-means clustering (**Fig. 3b,d**).

All methods except for *Cusanovich2018* and SnapATAC demonstrated declining performance with increased noise level (**Sup Fig. 2, 4a**). *Cusanovich2018* and SnapATAC were more robust to noise, showing no noticeable changes at increasing noise levels, while cisTopic was slightly more sensitive to noise; its performance dropped markedly when the noise level was increased to 0.4.

Next, the effect of the coverage on clustering performance was investigated. We progressively decreased the number of fragments per cell from a high coverage of 5,000 fragments, to a medium coverage of 2,500 fragments and 1,000 fragments, then to a low coverage of 500 fragments and finally to 250 fragments. The performance of all methods declined as coverage was decreased. (**Sup Fig. 4b, Sup Fig. 5, Sup Table 4-5-6-7-8**). *Cusanovich2018*, SnapATAC, Scasat, and Control-Naïve are relatively robust to low coverage and outperform other methods. cisTopic worked well with high coverage but in contrast to the above listed methods, was more sensitive to lower coverages (**Sup Fig. 5e**).

#### Simulated erythropoiesis datasets

Following the simulation of discrete sorted cell populations, we simulated three scATAC-seq datasets aimed at mimicking the continuous developmental erythropoiesis process and encompassing the following twelve populations: hematopoietic stem cells (HSC), common myeloid progenitors (CMP), megakaryocyte-erythroid progenitor (MEP), multipotent progenitors (MPP), myeloid progenitors (MyP), colony forming unit-erythroid (CFU-E), proerythroblasts (ProE1), proerythroblasts (ProE2), basophilic erythroblasts (BasoE), polychromatic erytrhoblasts (PolyE), orthochromatic erythroblasts (OrthoE) and OrthoE and reticulocytes (Orth/Ret). These datasets were generated as before with three noise levels (0, 0.2 and 0.4) and with 2,500 fragments per cell.

To first quantitatively evaluate the clustering solutions we used ARI, AMI and the homogeneity metrics (**Sup Fig. 6 and Sup Table 9)**. Without noise, SnapATAC, cisTopic BROCKMAN, *Cusanovich2018*, and Scasat consistently outperform the Control-Naïve across the three metrics (**Sup Fig.6a**). chromVAR as before, performs better using k-mers as features than when using motifs. SCRAT and scABC work as well as k-mers-based chromVAR. Again, methods such as Cicero and Gene Scoring that only summarize chromatin accessibility around TSS perform poorly. For scABC, Cicero and Gene Scoring, we also notice that there are significant discrepancies between the three clustering methods, but their performances become similar after PCA (scABC after PCA is equivalent to the Control-Naïve method). Again, we observe that PCA can significantly improve the clustering performance of Louvain for scABC, Cicero and Gene Scoring but not for chromVAR and SCRAT.

As before, to qualitatively assess population separation, we inspected UMAP projections applied to the noise-free simulated dataset (**Sup Fig. 6a**). In accordance with the quantitative comparison, cisTopic, *Cusanovich2018*, SnapATAC, and BROCKMAN demonstrate better performance in separating cell types compared to the Control-Naïve method and are able to further separate BasoE and PolyE. Moreover, SnapATAC can clearly distinguish CFU-E, ProE1, ProE2 while cisTopic, *Cusanovich2018*, and BROCKMAN are only able to separate ProE2 out of these three populations. Scasat performs similarly to the Control-Naïve method. chromVAR with k-mers as features and SCRAT are able to isolate six major groups including HSCs-MPPs, CMP, MEP, Myp, CFU-E-ProE1-ProE2, and BasoE-PolyE-OrthoE-Orth/Ret. chromVAR with k-mers performs well in preserving the order of CFU-E-ProE1-ProE2 and BasoE-PolyE-OrthoE-Orth/Ret. SCRAT can further separate BasoE-PolyE from OrthoE-Orth/Ret while mixing up CFU-E-ProE1-ProE2. As before, we noticed that chromVAR using k-mers as features obtained a better separation of cell types than when using motifs. scABC is able to preserve well the order of major groups in a continuous way but fails to separate CFU-E-ProE1-ProE2 and OrthoE-Orth/Ret. Cicero gene activity score and Gene Scoring mixed different cell types but after a simple PCA step they clearly separate cells into three major groups. scABC did not perform well and produced small noisy clusters with different cell types mixed together.

As expected, we observed that increasing the level of noise resulted in clustering performance decrease and a decline of visual separation of cell types for all the methods (**Sup Fig. 4c, Sup Fig. 6, Sup Table 10-11**). SnapATAC, cisTopic, and *Cusanovich2018* performed reasonably well when increasing the noise level, with SnapATAC the most robust among the three.

### Clustering performance on real datasets

Following the benchmark of the synthetic datasets, we assessed the performance of the methods on real datasets. These datasets were generated using different technologies: the Fluidigm C1 array[18], the 10X Genomics droplet based scATAC platform, and a recently-optimized split-pool protocol[1]. Each real dataset used was fundamentally different in its cellular makeup as well as size and subpopulation organization. Notably, as ‘true positive’ labels are not always available, in addition to the metrics used on the simulated datasets, here we introduced the RAGI, a simple metric based on the Gini Index that can be adopted when marker genes for the expected populations are known (**see Methods**). In our assessment of *Cusanovich2018*, to make a fair comparison, we use first the same set of peaks used for other methods instead of the peaks called from its pseudo-bulk-based procedure. However, since this strategy may be important for the final clustering performance, the pseudo-bulk based peak calling strategy is tested and discussed in a subsequent section.

#### Buenrostro2018 dataset

The first and smallest dataset we used in our benchmarking contains single cell ATAC-seq data from the human hematopoietic system (hereafter *Buenrostro2018*)[18]. This dataset consists of 2034 hematopoietic cells that were profiled and FACS-sorted from 10 cell populations including hematopoietic stem cells (HSCs), multipotent progenitors (MPPs), lymphoid-primed multipotent progenitors (LMPPs), common myeloid progenitors (CMPs) and granulocyte-macrophage progenitors (GMPs), GMP-like cells, megakaryocyte-erythroid progenitors (MEPs), common lymphoid progenitor (CLPs), monocytes (mono) and plasmacytoid dendritic cells (pDCs). **Fig. 4a** illustrates the roadmap of hematopoietic differentiation. For this dataset, the FACS-sorting labels are used as gold standard. The analysis details for each method are documented in **Sup Note 2.**

**Figure 4.**
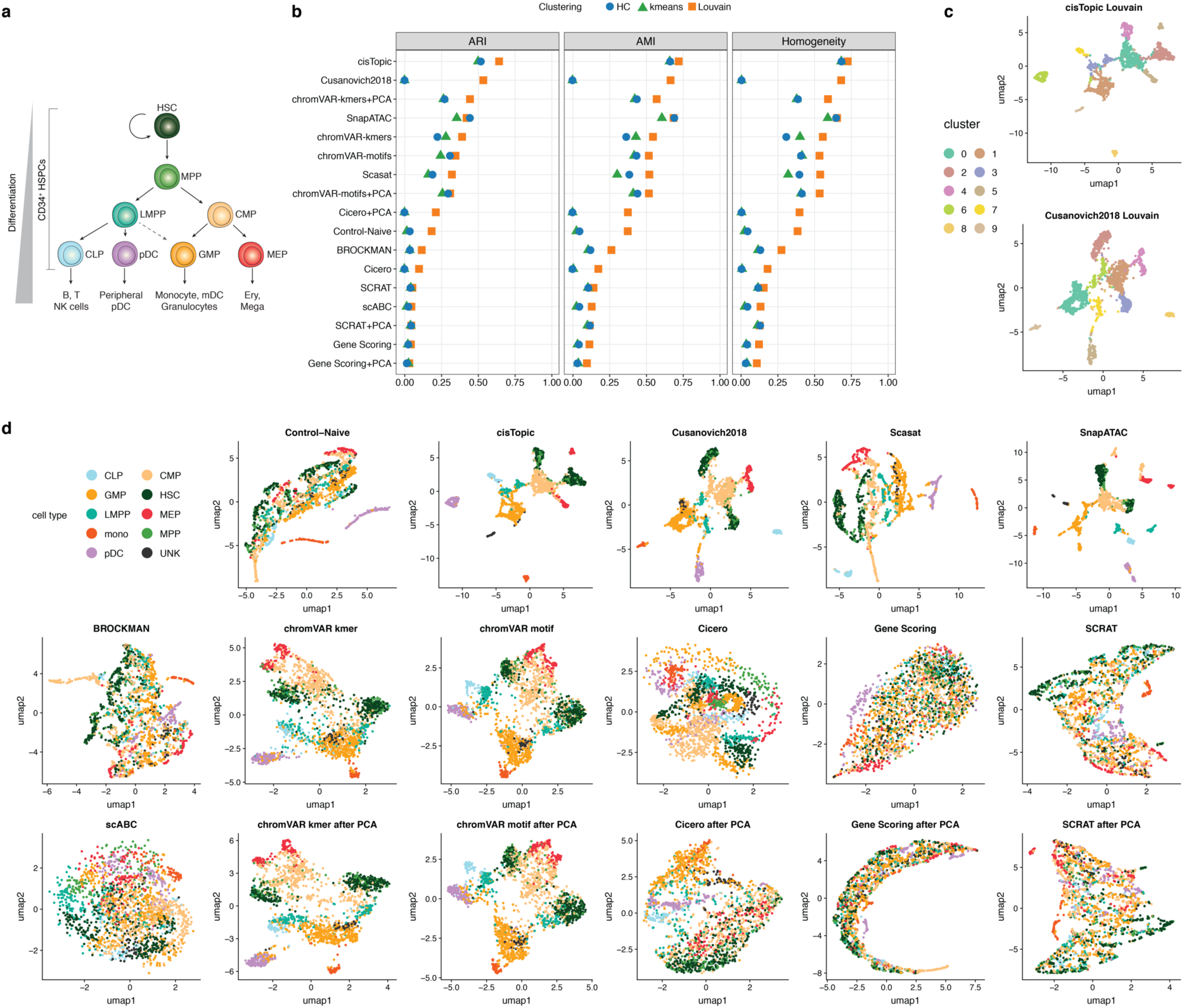
Benchmarking results using the *Buenrostro2018* scATAC-seq dataset. **(a)** Developmental roadmap of cell types analyzed. **(b)** Dot plot of scores for each metric to quantitatively measure the clustering performance of each method, sorted by maximum ARI score. **(c)** The two top-scoring pairings of scATAC-seq analysis method and clustering technique. UMAP visualization of the feature matrix produced by each method for the *Buenrostro2018* dataset. Individual cells are colored indicating the cell type labels shown in (a).

We started by evaluating the clustering solutions based on the feature matrices generated by the different methods. We used the same metrics used for the synthetic datasets: ARI, AMI and homogeneity (**Fig. 4b, Sup Table 12**). cisTopic, *Cusanovich2018*, chromVAR, SnapATAC, and Scasat outperform the other methods across all three metrics. We also observed that chromVAR with k-mers or TF motifs and with or without PCA performs consistently well. As before, k-mers-based features work better than motif-based features. This can be also observed when comparing BROCKMAN, another k-mers-based method, with SCRAT, which is a motifs-based method. TSS based methods including Cicero and Gene Scoring did not perform well. Cicero requires a preprocessing step to assess cell similarity; poor performance might be due to the internally incorrectly inferred coordinates (our assessment used the t-SNE procedure as suggested in their documentation). Implementing PCA consistently improves the performance of scABC (as mentioned before, scABC after PCA is equivalent to the Control-Naïve method) and Cicero but does not impact the performance of chromVAR, SCRAT, and Gene Scoring. We also observed that for this dataset, Louvain algorithm works consistently well across different metrics and methods and performs better than hierarchical clustering and k-means in almost all the cases.

We also qualitatively assessed the separation of different cell types by visualizing cells in UMAP projections based on the FACS-sorted labels (**Fig. 4d**) and clustering solutions (**Sup Fig. 7**). **Fig. 4c** shows the best two combinations based on ARI: cisTopic with Louvain and *Cusanovich2018* with Louvain (the complete ranking is presented in **Sup Table 12**).

As **Fig. 4d** shows, in accordance with the clustering analyses, cisTopic, *Cusanovich2018*, Scasat, SnapATAC, and chromVAR can generally separate cell types, and reasonably capture the expected hematopoietic hierarchy. cisTopic and SnapATAC show a clear and compact separation among groups, with SnapATAC recovering finer structure within each cell type cluster. chromVAR with k-mers or motifs corresponds to a more continuous progression of the different cell types. Control-Naïve and BROCKMAN perform comparably in distinguishing cell types and preserving the continuous hematopoietic differentiation. Cicero gene activity scores, SCRAT, and scABC show ambiguous patterns of distinct cell populations while Gene Scoring fails to separate different cell types. For Cicero gene activity score, after performing PCA, the separation of different cells is noticeably improved. For SCRAT, performing PCA does not show clear improvement.

#### Peripheral blood mono nuclear cells (PBMCs) 10X dataset

Next, we investigated a recent dataset produced by 10X Genomics profiling peripheral blood mononuclear cells (PBMCs) from a single healthy donor. In this dataset, 5335 single nuclei were profiled (~42k read pairs per cell); no cell annotations are provided. Based on recent studies [9, 19], we expected ~8 populations: CD34+, Natural Killer and Dendritic cells, Monocytes, lymphocyte B and lymphocyte T cells, together with terminally differentiated CD4 and CD8 cells. Therefore, we used 8 as the number of expected populations for the clustering procedures. The analysis details for each method are documented in **Sup Note 3**.

Several marker genes have been proposed to label the different populations or to annotate clustering solutions for PMBCs [9, 19]. To measure cluster relevance based on these marker genes, we can annotate the clusters (or alternatively any group of cells) according to the accessibility values at those marker genes. In addition, accessibility at marker genes should be more variable between clusters than accessibility at housekeeping genes (since they should be, by definition, more equally expressed across different populations). Based on these ideas, we proposed and calculated the Residual Average Gini Index (RAGI) score (see **Methods**) contrasting marker and housekeeping genes (**Fig. 5a, Sup Table 13**). For reasonable clustering solutions, we expect that the accessibility of marker genes defines clear populations corresponding to one or few clusters, whereas accessibility of the housekeeping genes is broadly distributed across all the clusters.

**Figure 5.**
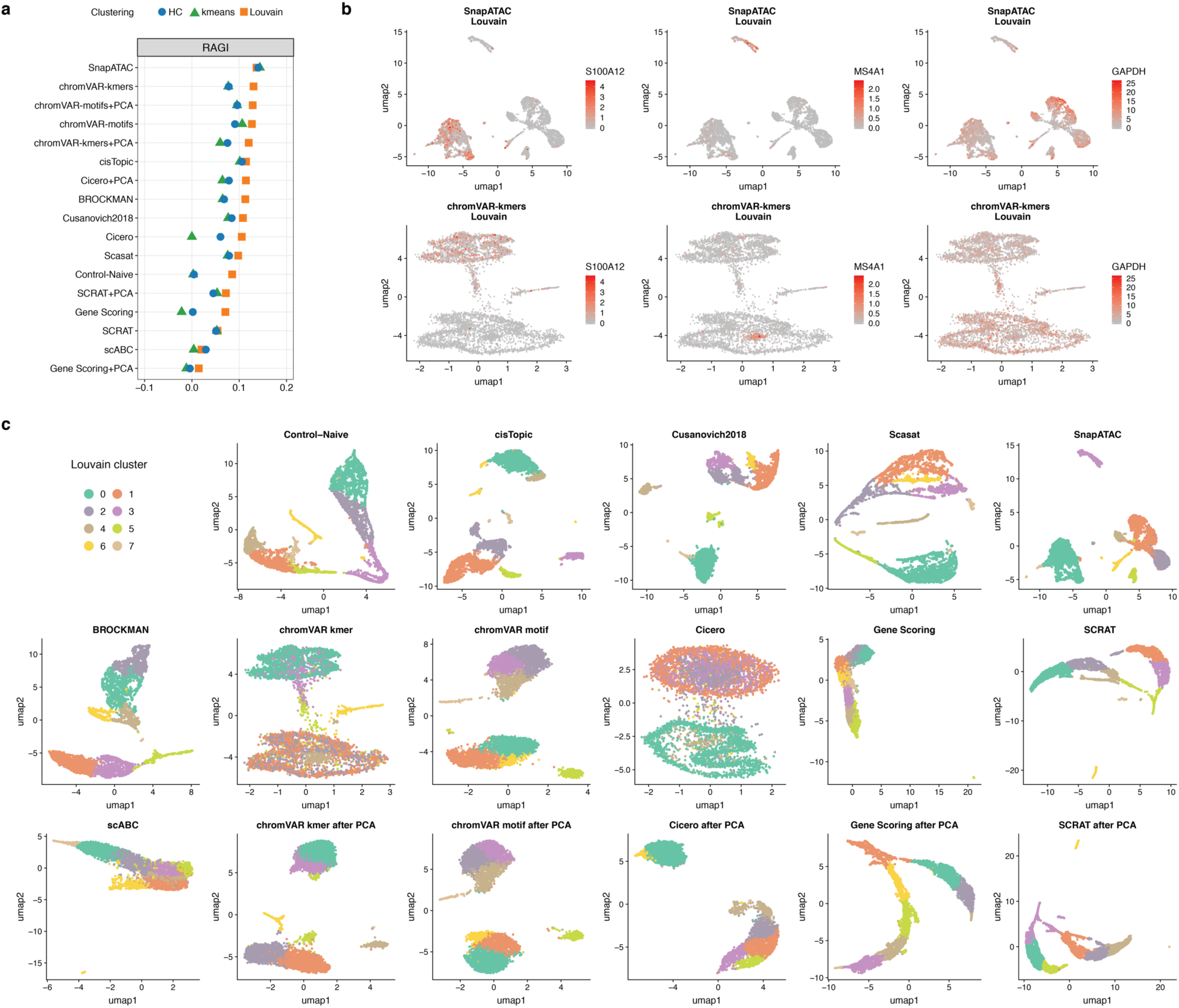
Benchmarking results using scATAC-seq data for 5k Peripheral blood mononuclear cells (PBMCs) from 10x Genomics. **(a)** Dot plot of RAGI scores for each method, sorted by the maximum RAGI score. A positive RAGI value indicates that a method is able to produce a clustering of PBMCs in which chromatin accessibility of each marker gene is high in only a few clusters relative to the number of clusters with high accessibility of housekeeping genes. **(b)** UMAP visualization of the feature matrix produced by the top two methods (top row: SnapATAC, bottom row: chromVAR using kmers). Chromatin accessibility of S100A12 (left, Monocyte marker gene), MS4A1 (center, B-cell marker gene) and GPDH (right, housekeeping gene) are projected onto the visualization. **(c)** UMAP visualization of the feature matrix produced by each method for the 5k PBMCs dataset from 10x genomics. Individual cells are colored indicating cluster assignments using Louvain clustering.

As expected, methods with the highest performance such as SnapATAC and chromVAR, showed a higher average accessibility for just one cluster for the same marker gene, while lower performing methods such as SCRAT or Gene Scoring showed higher average accessibility in multiple clusters for the same marker gene, further motivating the use of the RAGI metric (**Sup Fig. 8**). **Fig. 5b** shows for the top two performing methods based on RAGI (SnapATAC and chromVAR with k-mers) the gene accessibility patterns for 3 genes (S100A12 - Monocytes-specific, MS4A1 - B cells specific and GAPDH - housekeeping.) The same three genes are also shown in UMAP plots of the other methods (**Sup Fig. 9**). Again, we observed that Louvain algorithm performed better than k-means and hierarchical clustering for almost all scATAC-seq methods. Importantly, negative RAGI score for a method (see for example the solutions obtained by the Gene Scoring **in Fig. 5a, Sup Fig. 9**) may suggest that its clustering solutions are defined by housekeeping genes rather than informative marker genes

We also qualitatively evaluated the clustering solutions of the different methods using UMAP projections (**Fig. 5c, Sup Fig. 10**). We observed two major groups for all methods except for scABC. Among these methods, the UMAP projections based on feature matrices obtained by Control-Naïve, cisTopic, *Cusanovich2018*, Scasat SnapATAC, BROCKMAN and chromVAR showed additional smaller groups and finer structures. For Cicero gene activity scores, performing PCA helps to improve the separation of more putative cell types. Instead for SCRAT and Gene Scoring, the PCA step did not improve the separation.

Given that the ranking of methods in datasets with ground truth is similar to the ranking based on the RAGI metric, we believe this simple approach is a reasonable surrogate metric that can be useful for evaluating unannotated datasets, a common scenario in single cell omics studies.

#### sci-ATAC-seq mouse dataset

The last dataset analyzed in our benchmark consists of sciATAC-seq data from 13 adult mouse tissues (bone marrow, cerebellum, heart, kidney, large intestine, liver, lung, pre-frontal cortex, small intestine, spleen, testes, thymus and whole brain), of which 4 were analyzed in duplicate for a total of 17 samples and 81,173 single cells[1]. Each tissue can be interpreted as a coarse ground truth, used later to evaluate clustering solutions (**Fig. 6a**). The analysis details for each method are documented in **Sup Note 4**.

**Figure 6.**
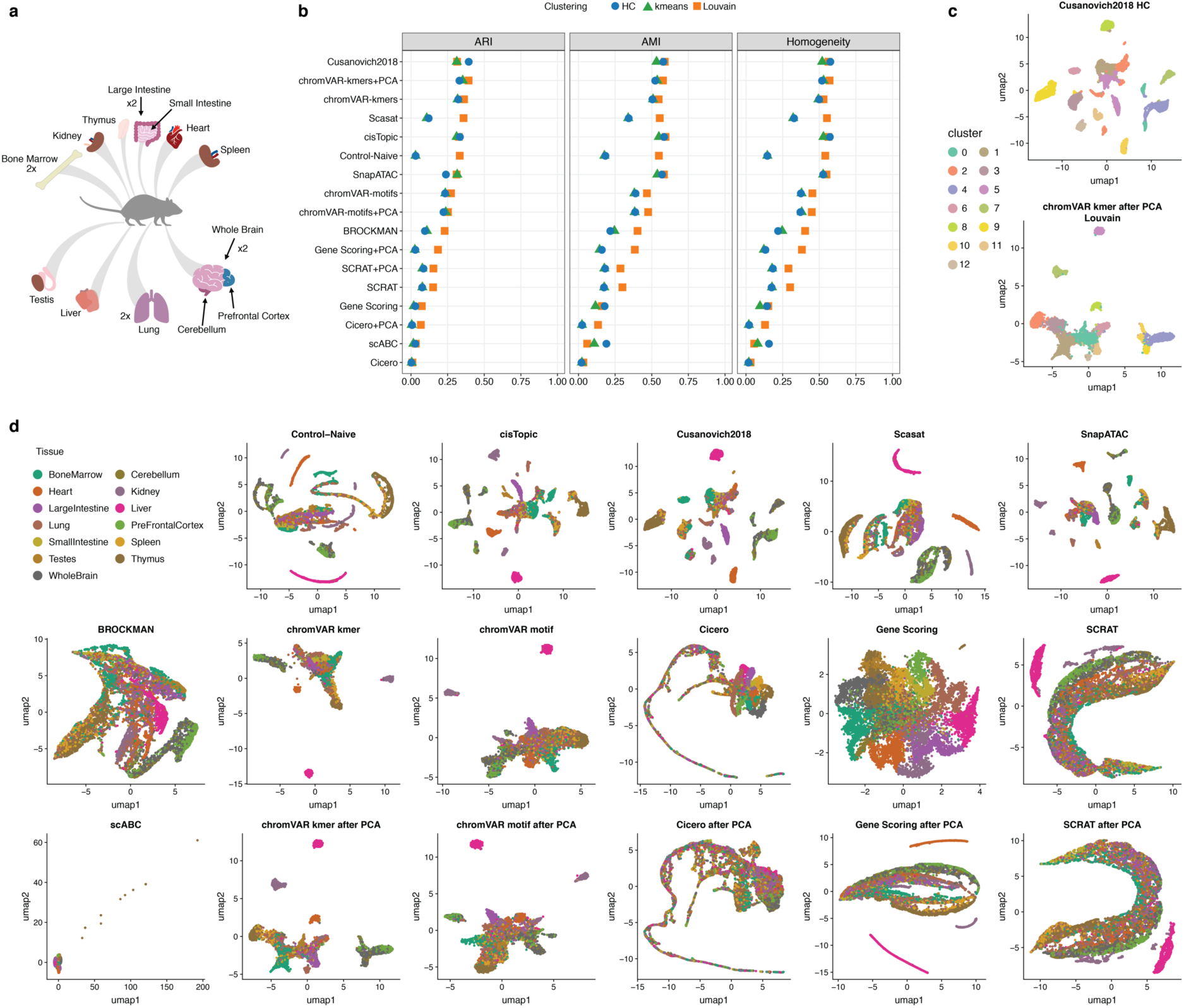
Benchmarking results using the downsampled sci-ATAC-seq mouse dataset from 13 adult mouse tissues. **(b)** Dot plot of scores for each metric to quantitatively measure the clustering performance of each method, sorted by maximum ARI score. **(c)** The two top-scoring pairings of scATAC-seq analysis method and clustering technique. Cell cluster assignments from each method are shown using the colors in the legend on the left. **(d)** UMAP visualization of the feature matrix produced by each method for the downsampled sci-ATAC-seq mouse dataset. Individual cells colors indicate the cell type.

Despite using a machine with 1 TB of memory, almost all the methods failed to even load this dataset, owing to its size. The only method capable of processing this dataset in a reasonable time was SnapATAC (~700 minutes). The other methods failed to run due to memory requirements. To understand the causes of this failure we did an in-depth analysis of their scalability looking at their source code (**Sup Note 5**). Briefly, we found that the majority of the methods try to load the entire dataset in the central memory while SnapATAC uses a custom file format (.snap) based on HDF5 (https://support.hdfgroup.org/HDF5/whatishdf5.html), allowing out of core computation by efficiently and progressively loading in the central memory only the data chunks required at any given moment of the analysis.

On this dataset, SnapATAC was able to correctly cluster cells of the following tissues: kidney, lung, heart, cerebellum, whole brain and thymus. However, for the other tissues, including bone marrow and small intestine, cells are distributed in groups of mixed cell types (**Sup Fig.11**), as reflected by the score of the three metrics used for the other datasets evaluation (**Sup Table 14**), i.e. ARI= (HC=0.24, k-means=0.34, Louvain=0.39), AMI=(HC=0.55, k-means=0.55, Louvain=0.62), Homogeneity=(HC=0.52, k-means=0.54, Louvain=0.60).

To gain insight on the performance of the other methods on this this dataset, we randomly selected 15% of cells from each sample to construct a smaller sciATAC-seq dataset consisting of 12,178 cells.

As **Fig. 6b** shows *Cusanovich2018*, k-mer-based chromVAR, cisTopic, SnapATAC, Scasat and Control-Naïve perform comparably well and have noticeably better clustering scores than the other methods (**Sup Table 15**). Consistent with what we observed previously, peaks or bins level methods generally work better. In this dataset, k-mers-based chromVAR and its combination with PCA transformation performs equally well as peaks or bins-level methods and better than the motifs-based methods. Simply counting reads within peaks (scABC) and gene-level-featurization-based methods (Gene Scoring and Cicero) perform poorly overall. Adding a PCA step improves noticeably scABC (scABC after PCA is the same as Control-Naïve) and Gene Scoring. It also slightly improves Cicero but it does not affect chromVAR and SCRAT.

As before, all the clustering solutions of the different methods were visualized in UMAP plots (**Sup Fig. 12**). The top two combinations, i.e. *Cusanovich2018* and chromVAR k-mers with PCA, are visualized in **Fig.6c**. To visually compare the separation of the different tissues across methods, we also inspected UMAP plots where cells are colored based on the tissue of origin. Similar to what we observed using the clustering analysis, cisTopic, *Cusanovich2018*, and SnapATAC are able to separate cells into the major tissues and also to capture finer discrete groups. The Control-Naïve method and Scasat are also able to distinguish the major tissues but show some mixing within each discrete cell population. K-mer-based chromVAR can separate out liver, kidney, and heart tissues and present the other tissues within a continuous bulk population while preserving the structure of the distinct tissues. We observed that after running PCA, k-mer-based chromVAR can recover an additional group of cells within the lung tissue and also detect finer structure within the cells from the brain. Compared with k-mer-based features, motif-based chromVAR and its combination with PCA transformation distinguished fewer tissue groups while mixing more cells from different tissues. BROCKMAN recovered a continuous structure with the different tissues but does not distinguished them clearly. Similarly, Gene Scoring put cells from different tissues into a big bulk population with limited separation. PCA improved its ability to separate out a few tissues, including liver, heart, and kidney. SCRAT and Cicero gene activity scores mixed most of the cells from different tissues and performed poorly on this dataset with or without PCA.

### Clustering performance summary

To assess and compare the overall performance of scATAC-seq analysis methods, we ranked the methods based on each metric (ARI, AMI, Homogeneity, RAGI) by taking the best clustering solution for the three real datasets (*Buenrostro2018* dataset, *PBMCs 10X* dataset, and the down-sampled *sci-ATAC-seq mouse* dataset) and two synthetic datasets (simulated bone marrow dataset and simulated erythropoiesis dataset with the moderate noise level of 0.2 and a medium coverage of 2500 fragments per cell). Then for each dataset except for the *PBMCs 10X* dataset, we calculated the average rank across ARI, AMI, and Homogeneity. For the *PBMCs 10X* dataset, RAGI is calculated instead (**Sup Fig.13a**). Lastly, we calculated the average rank across different datasets. According to the average ranking, SnapATAC, cisTopic and *Cusanovich2018* are the top three methods to create feature matrices that can be used to cluster single cells into biologically-relevant subpopulations (**Fig. 7a**). SnapATAC consistently performed well across all datasets. Both cisTopic and *Cusanovich2018* demonstrated satisfactory performance across all datasets except for the 10X PBMCs dataset.

**Figure 7.**
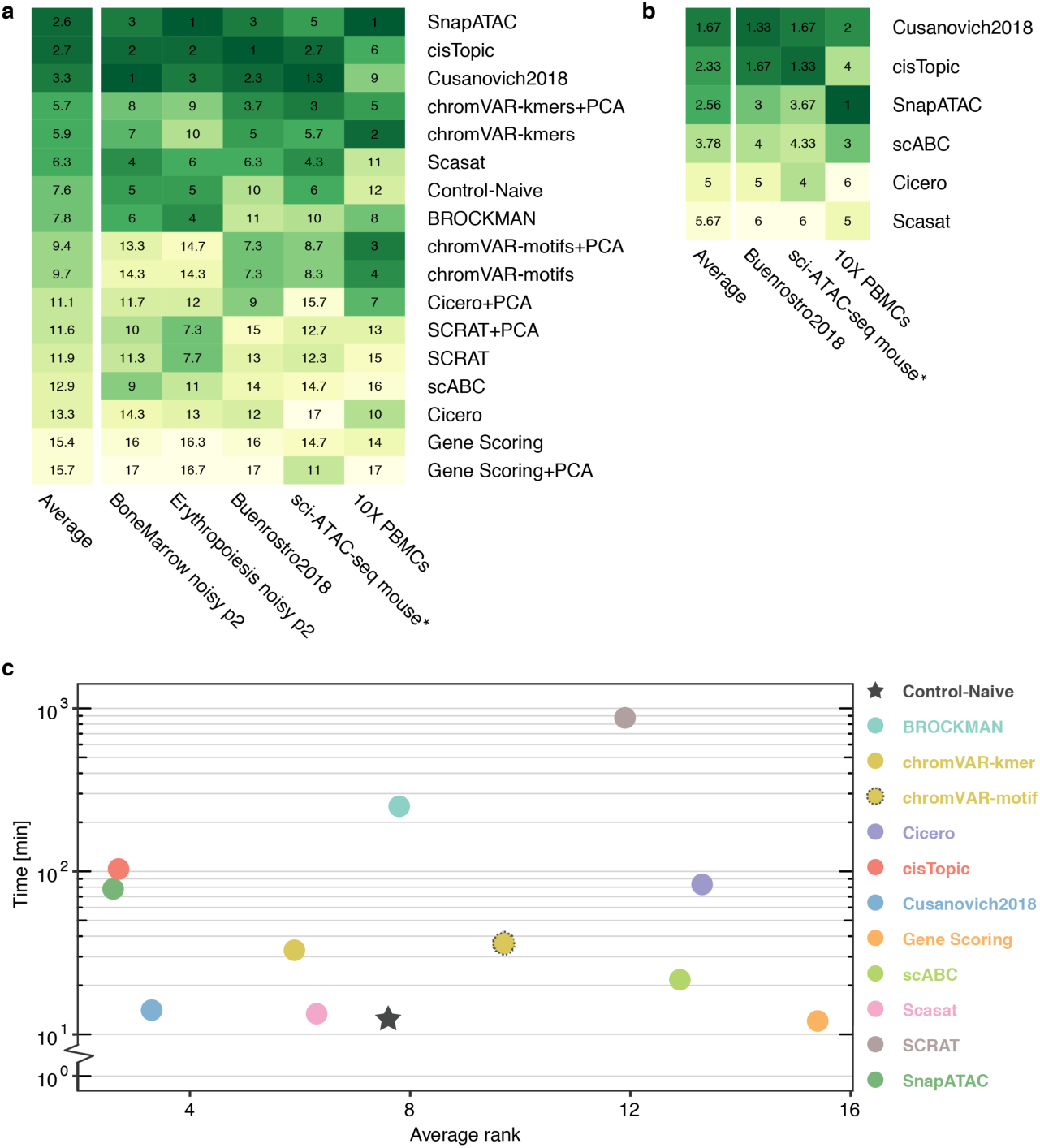
Aggregate benchmark results. **(a)** For each method, the rank based on the best-performing clustering method is measured for each metric (e.g. ARI, AMI, H, or RAGI). The average metric ranks for each dataset were used to calculate a performance score for each method. Each method was then assigned a cumulative average score based on its performance across all datasets. * indicates a downsampled dataset of the indicated original dataset. **(b)** For methods that specify an end-to-end clustering pipeline, average rank and cumulative average scores for each method were calculated as in (a). **(c)** Plot of running time against performance for each method. Cumulative average scores, which were calculated in part (a) are shown on the x-axis, and the average running time across the three real datasets (*Buenrostro2018*, 10X PBMCs, and downsampled sci-ATAC-seq mouse) is shown on the y-axis.

Generally, methods that implement a dimensionality reduction step work better (SnapATAC, cisTopic, *Cusanovich2018*, Scasat, Control-Naïve, and BROCKMAN) than those without it (SCRAT, scABC, Cicero, and Gene Scoring). We also observed that chromVAR performs better in real datasets than in simulated datasets and that the kmer-based version of chromVAR consistently outperforms motif-based chromVAR. For the methods that do not implement dimensionality reduction, the PCA step does not always improve the performance except for scABC and Cicero, in which the PCA transformation consistently boosts the results. Interestingly, we observed that regardless of the method, the PCA consistently improves the clustering solutions obtained by the Louvain algorithm.

#### Keeping the first PC vs removing the first PC

In preparing this manuscript, we noticed that in some cases, the first principal component (PC) may only capture variation in sequencing depth instead of biologically meaningful variability. To make a thorough assessment of how the first PC affects the clustering results, we compared the effect of keeping vs removing the first PC on the three real datasets (for this comparison we consider both the methods that implemented PCA and the combination of PCA and the methods that did not implement a dimensionality reduction step) (**Sup Fig.14**). Across all three datasets, we observe that for Control-Naïve, BROCKMAN, SCRAT-PCA, and Gene Scoring-PCA, removing the first PC consistently helped in better separating the different populations in UMAP projections and improved clustering performance. In contrast, the performance of chromVAR-PCA with motifs as features consistently dropped after removing the first PC. *Cusanovich2018* and SnapATAC performed similarly before and after removing the first PC across all datasets. For Cicero-PCA, removing first PC did not clearly affect its performance in *Buenrostro2018* and 10X PBMCs datasets but improved its performance in the down-sampled sci-ATAC mouse dataset.

Generally, the methods that implement binarization (e.g. *Cusanovich2018*, SnapATAC) or that implement cell coverage bias correction (e.g. chromVAR, SnapATAC), tend to be less affected by the sample sequencing depths. Therefore, for these methods we believe that the first PC does not capture the library size and removing it does not help to improve the clustering results. On the contrary, for methods that do not implement any specific step to correct for potential artifacts associated with sequencing depth, the first PC is more likely to capture biologically irrelevant factors and therefore may reduce biology-driven differences. However, this operation must be applied with caution, since removing the first component could also in some cases remove some biological variation (e.g. motif-based chromVAR).

#### Clustering performance when running methods as end to end pipelines

When designing this study, we reasoned that a benchmark procedure could be approached from two very different perspectives. The first is the end user perspective, i.e. a user that runs a method as a black box following the provided documentation with the goal to obtain a reasonable clustering solution without worrying too much about the internal design choices and procedures. In these settings, it is not trivial to systematically compare the methods and understand which part related to the featurization may influence the final clustering performance, especially if also the clustering algorithms used are different. The second perspective that was used instead in the rest of this benchmarking effort is the developer perspective, i.e. we tried to understand what are the key steps of each method that can boost clustering performance of common clustering approaches. Regardless, we reasoned that it is important to provide some insights on the user perspective, since some readers will use the tested methods as end-to-end pipelines. Therefore, we also compared the clustering solutions produced by running the complete analysis pipelines as outlined in tutorials for the methods that explicitly implement a clustering step (see **Sup Note 6**). We evaluated the clustering results using ARI, AMI and Homogeneity for the *Buenrostro2018* and sci-ATAC-seq mouse datasets, and RAGI for the PBMCs 10X dataset (**Sup Table 16-17-18**). We observe the top three methods, i.e. Cusanovich2018, cisTopic and SnapATAC, still outperform the other methods but with a slightly different ranking. (Cusanovich2018 is ranked first followed by cisTopic and SnapATAC, Fig. 7b, Sup Fig. 13b). Also, both scABC and Cicero performed better than Scasat in this analysis. Interestingly, we observed that SnapATAC, cisTopic, Cusanovich2018, and Scasat have even better clustering solutions in our benchmarking framework compared to using their own clustering approach. On the other hand, scABC and Cicero had better clustering results when running their own clustering procedure. scABC uses an unsupervised clustering method tailored to single cell epigenomic data (including scATAC-seq). Although it uses the naïve peaks-by-cells raw count as its feature matrix, it calculates cells weights by considering their sequencing coverage and giving more weight to cells with higher number of reads. Also, it performs two steps of clustering by using weighted k-medoid algorithm based on Spearman rank correlation to find landmarks first and then assigns cells to the landmarks. These specific steps help improve its clustering performance. For the Cicero clustering workflow, we used the gene activity scores and, as proposed in their tutorial, functions from Monocle2, to (i) normalize the scores and (ii) reduce the dimensionality with tSNE by using the top PCs before clustering cells. These extra steps helped in improving its clustering solutions. This suggests that appropriate normalization steps need to be properly performed to improve clustering analysis, in addition to simple transformations like binarizing counts and/or performing a PCA.

Taken together, based on these analyses, we recommend using SnapATAC, cisTopic, or *Cusanovich2018* to cluster cells in meaningful subpopulations. This step can be followed by methods such as Cicero, Gene Scores or with TF motifs (e.g. chromVar) to annotate clusters and to determine cell types in an integrative approach.

### Important considerations in defining informative regions for scATAC-seq analyses

Feature sets of informative peaks for scATAC analyses may be computed from bulk samples available through large scale consortia such as ENCODE[20] and ROADMAP[21] or more precise tissue-specific cell types as in the murine ImmGen Project[22]. However, scATAC-seq analyses often require *de novo* inference of dataset-specific accessibility peaks in order to resolve cell types and regulatory activity.

To date, there are three major methods for generating peak sets for scATAC experiments. The first strategy (pseudo-bulk from all single cells, PB-All) for inferring peaks is to call peaks on a pseudo-bulk sample omposed of all the reads from all cells in the library. The second (pseudo-bulk from FACS, PB-FACS) is to call peaks in *a priori*-defined cell types isolated by FACS-sorting. A consensus peak set can be defined by combining summits of individual peaks using an iterative algorithm [5, 18, 23]. Finally, a third strategy (pseudo-bulk from clades, PB-Clades) uses a pre-clustering of cells to define initial populations[1, 10]. Subsequent peak calling is performed in each initial cluster. Aggregate peak sets can then be defined from synthesizing the summits of each cluster-specific peak set as described above.

#### Bulk ATAC-seq peaks vs aggregated scATAC-seq peaks

To evaluate the effect of using peaks obtained from bulk ATAC-seq data versus peaks obtained from aggregated single cell profiles, we reanalyzed the *Buenrostro2018* dataset in which both are available (**Sup Fig. 15–16**). Here we considered only the methods that use peaks as input (i.e. SnapATAC, SCRAT, BROCKMAN are excluded). For the aggregated scATAC-seq peaks, we merged cells of the same cell type based on the FACS sorting labels and performed peak calling within each cell type. Then peaks defined within each cell type were merged. For most methods we did not observe clear differences in performance between the two input peak strategies. For cisTopic, *Cusanovich2018*, and Cicero, aggregated scATAC-seq peaks overall perform better across all three metrics (**Sup Fig. 17a, Sup Table 19**).

We also tested the strategy of defining pseudo bulk samples from clades when no sorting labels are provided. *Cusanovich2018* is the only method that provides a workflow to identify initial clades and call peaks within each clade. It counts reads within the fixed-size windows and pre-clusters cells using hierarchical clustering to define initial clades from which peaks are called. We applied this strategy to all three real datasets (**Sup Fig. 18**). We observed that in all three datasets, *Cusanovich2018* performs well in identifying the isolated major groups and the identified clades match well the labels provided, including FACS-sorted labels, cell-ranger clustering solutions, and known tissues labels. Overall the *Cusanovich2018* ‘pseudo bulk’ strategy for defining *de novo* peaks is able to capture the heterogeneity within single cell populations and can serve as a promising unsupervised way to define pseudo bulk subpopulations and to perform peak calling.

#### The effect of excluding regions using the ENCODE blacklist annotation

cisTopic, Scasat, SCRAT, and SnapATAC employ a blacklist filtering step to remove features annotated by ENCODE as belonging to a subset of genomic regions, which harbor the potential to produce artifacts in downstream analysis steps [24]. cisTopic and Scasat perform a peak filtering in the pre-processing steps of their pipeline. Our benchmarking pipeline makes use of the ENCODE ATAC-seq pre-processing pipeline, which removes peaks overlapping with regions on the blacklist annotations list. Therefore, we tested the remaining two methods, which do not use peaks as features, SCRAT or SnapATAC. In particular, we wanted to test whether we would observe any change in downstream clustering performance upon opting to perform a blacklist removal step. Through a qualitative and quantitative comparison of clustering performance, we determined that methods, which remove features according to blacklist annotations show no considerable advantage over those that permitted such features (**Sup Fig.19**).

#### Rare cell type-specific peak detection

As all cell identities may not be pre-defined in complex tissue types, we sought to examine PB-All and PB-Clades strategies to infer a chromatin accessibility feature set from the scATAC-seq libraries directly. To achieve this, we established a simulation setting where we mixed bulk ATAC-seq data from three sorted populations (B-cells, CD4+ T-cells, and monocytes from the PBMCs 10X dataset) that would be mixed in complex tissue (i.e. peripheral blood mononuclear cells) (**Sup Fig. 17b**). After peak calling on both the synthetic bulk and isolated reads from each cell type, we inferred the proportion of cell type-specific peaks from the minor cell population that were captured by the peak calling in the synthetic bulk mixture (see **Methods**).

Overall, the results indicate that cell type-specific peaks may be vastly underestimated from performing peak calling on the mixture of single cells (PB-All) (**Sup Fig. 17b**). Specifically, only ~18% of cell type-specific peaks from very rare (1% prevalence) or ~40% from rare (5% prevalence) cell populations were detected when peaks were called when treating the heterogenous source as a synthetic bulk experiment. Consequently, as these peaks would be vastly under-represented in a consensus peak set, virtually all computational algorithms will fail to identify rare populations. Moreover, as many common quality-control measures for scATAC involve filtering based on the proportion of reads in peaks, these cell populations may be under-represented in quality-controlled datasets.

As observed in other studies [1, 25], these results suggest calling peaks on PB-All may result in sub-optimal performance. Alternatively, when isolated populations have been profiled (for example by FACS) peak sets can be defined by calling peaks using data from cells in each pre-defined population separately as discussed in the previous section since this enables the resolution of rare subpopulations (for example HSC in the hematopoietic system).

#### Frequency-based peak selection vs intensity-based peak selection

*Cusanovich2018* selects peaks that are present in at least a specified percentage of cells before performing TF-IDF transformation, while scABC selects peaks with the most reads to cluster cells. To evaluate the effect of selecting peaks based on their representation in the cell population or based on their intensity (defined as the sum of reads in that peak in all samples), we focus on the two methods that implement the step of peak selection, *Cusanovich2018* and Control-Naïve (equivalent to scABC+PCA).

To assess the two peak selection strategies, we ran both *Cusanovich2018* and Control-Naïve on both simulated bone marrow dataset at noise level of 0.2 with a coverage of 2500 fragments and the *Buenrostro2018* dataset by varying the cutoffs for peak inclusion (**Sup Fig. 20–21**). We calculated the intensity of peaks by counting the number of reads across all cells and calculated the frequency of peaks by counting the number of cells in which a peak is observed. For this analysis we selected the top peaks based on intensity and frequency with the following cutoffs: top 100%, 80%, 60%, 40%, 20%, 10%, 8%, 6%, 4%, %2, 1%.

For both *Cusanovich2018* and Control-Naïve, the two peak selection strategies have similar clustering result scores when varying the cutoff (**Sup Fig. 20a-b,21a-b**). We observed reasonable and stable clustering performance using more than 20% of the ranked peaks. As the number of peaks is reduced, the scores start to decline noticeably and decrease almost monotonically. Below 1%, both methods perform poorly. In addition, we observed that the Louvain method produces more stable results than hierarchical clustering and k-means across the considered settings.

### Running time of different methods

In our analysis, we also collected the running time of each method on both simulated and real datasets (see **Sup Note 6**). For the simulated datasets, we only reported the execution time necessary to build a feature matrix starting from a peaks-by-cells count matrix. For real datasets, we considered the execution time to build a feature matrix from bam files. The running times are shown in **Sup Fig. 22 (Sup Table 20)**. All the tests were run on a machine with an Intel Xeon E5-2600 v4 X CPU with 44 cores and 1 TB of RAM with the CentOS 7 operating system. When analyzing real datasets with methods that rely on peaks but do not provide an explicit function to construct a peaks-by-cells matrix (*Cusanovich2018*, Cicero, Gene Scoring and Scasat), we ran the same script on a Linux cluster to obtain the peaks-by-cells matrix such that the execution time of this step is equivalent across these methods. It is worthwhile to mention that not all the methods of this benchmark support parallel computing. For the methods that support parallel computing, including SnapATAC, chromVAR, and cisTopic, the execution time was reported using 10 cores. For the rest of methods, we run them using a single core. We selected this number reasoning that a typical lab may not have access to a machine with 44 cores and instead may use a mid-size computing node with 8-12 cores. Notably, SnapATAC was the only method capable of processing the full sci-ATAC-seq mouse dataset (~80,000 single cells).

As shown in **Sup Fig. 22**, BROCKMAN and SCRAT have the largest greater execution time in all the real datasets while the methods that use a custom script to obtain a peaks-by-cells matrix tend to have shorter execution time (e.g. Scasat, *Cusanovich2018*, Gene Scoring).

We also assessed the scalability of methods with respect to the increasing coverage (250, 500, 1000, 2500 and 5000 fragments per peaks). We observe that with the increase of read coverage, for cisTopic there is an exponential increase of the running time whereas for other methods, the running time stays stable or increases linearly (**Sup Fig. 22, Sup table 21**).

Finally, we compared execution time vs clustering performance (**Fig. 7c**). Interestingly, the most accurate methods (SnapATAC, cisTopic and Cusanovich2018) have a reasonable running time while outperforming the other methods for clustering quality across all the datasets. Considering the computational time as an important factor that must be carefully evaluated before the implementation of any bioinformatics pipeline, we believe that *Cusanovich2018* is the best in balancing clustering performance with execution time.

## Discussion

scATAC technologies enable the epigenetic profiling of thousands of single cells, and many computational methods have been developed to analyze and interpret this data. However, the sparsity of scATAC-seq datasets provides unique challenges that must be addressed in order to perform essential analyses such as cluster identification, visualization and trajectory inference [26, 27]. Moreover, the rapid technological innovations that facilitate profiling accessible chromatin landscapes of 10^4^ or 10^5^ cells provide additional computational challenges to efficiently store and analyze data.

In this study, we compared ten computational methods developed to construct informative feature matrices for the downstream analysis of scATAC-seq data. We developed a uniform processing framework that ranks methods based on their ability to discriminate cell types when combined with three common unsupervised clustering approaches, followed by evaluation of three well-accepted clustering metrics. We evaluated these methods on thirteen datasets, three of those obtained using different technologies (Fluidigm C1, 10X, and sci-ATAC), and five consisting of simulated data with varying noise levels. These datasets comprise cells from different tissues in both mouse and human.

In addition to identifying various methodologies that perform optimally on real and simulated data, our benchmarking examination of scATAC-seq methodologies reveals general principles that will inform the development of future algorithms. First, peak-level or bin-level feature counting generally performs better in distinguishing different cell types followed in turn by k-mer-level, TF motifs-level, and gene-centric level summarization. We interpret this finding as an indication of the complexity of gene regulatory circuits where precise enhancer elements may have distinct functions that cannot be sufficiently approximated by sequence context or proximity to gene bodies alone. Second, we note that the methods that implement a dimensionality reduction step generally perform better in the separation of cell types, since this step may help to remove the redundancy between a large number of raw features and to mitigate the effect of noise. Third, for the methods that do not implement a dimensionality reduction step, simply adding a PCA step could significantly improve the clustering results. In fact, PCA generally boosts Louvain clustering results. For methods that do not account for the differing sequencing coverage of cells, the first PC could be used to capture and correct for sample depth differences. In this case, removing the first PC may improve the performance of these methods. Fourth, we observe that the Louvain method overall performs more consistently and accurately than k-means and hierarchical clustering. In contrast, k-means and hierarchical clustering are more sensitive to outliers and may result in suboptimal clustering solutions since some of clusters may correspond to single or few outlier cells. Fifth, the robustness of different methods to noise and coverage varies among different datasets. Among the top three methods, cisTopic is the most penalized by low coverage. Sixth, it was also observed that inappropriate transformations, such as log2 transformation and normalization based on region size as implemented in SCRAT may impact negatively clustering performance.

We observe that many methods fail to scale to larger datasets, which are now available due to improvements in split-pool technology and droplet microfluidics. As technologies improve and individual labs and international consortia lead efforts to generate ever larger single-cell datasets, scalability will be an unavoidable goal of method developments on a par with accuracy. As many of our evaluated methods were designed in the context of data generated from the Fluidigm C1 platform (which produces ~10^2^ cells), such approaches were often incapable of analyzing large datasets. In particular, the sci-ATAC-seq mouse dataset served as a useful resource to test the scalability of the methods that were benchmarked (~80,000 cells). Notably, our evaluation demonstrates that only SnapATAC was able to scale to process and analyze this large dataset. Future methods must be capable of processing datasets of this size especially adopting efficient data structures that allow out of core computing. Our findings reinforce the need for methods that not only are accurate but highly scalable for scATAC-seq data processing.

Defining regions is an important step in constructing feature matrices. Selecting informative regions generally improves downstream analyses such as clustering to capture heterogeneity within cell populations. Peak calling is a popular and straightforward way to define regions of interest. We observe that clustering performance is not generally impacted by using peaks defined from bulk ATAC-seq data vs using peaks obtained from aggregating single cell data based on FACS-sorting labels. However, performing peak calling by simply pooling reads from single cells may obfuscate peaks specific to rare cell populations leading to failures in uncovering them. In addition, the *Cusanovich2018* approach to identify pseudo-bulk clades is a promising unsupervised way to perform *in silico*-sorting without relying on FACS-sorting labels. This strategy potentially serves as a suitable way to preserve peaks specific to rare cell types. Also choosing an appropriate number of peaks is important for improving the downstream analysis (for example based on intensity/frequency-based given that they perform similarly).

We are aware of current limitations in our benchmarking effort. We have compared single cell ATAC-seq methods based on their ability to separate discrete cell populations; however, this might not be ideal when dealing with a continuous cell lineage landscape. We observe that chromVAR generally works better in preserving a continuous space while SnapATAC tends to break a putative landscape into discrete populations. The choice of method is ultimately case-specific and may be driven by the downstream application. For example, the feature matrix obtained by chromVAR may be more suitable for trajectory inference [26] while the one obtained from SnapATAC may be more appropriate better identify discrete and well separated cell populations by clustering. We acknowledge also that not all tested methods were specifically designed to produce clustering results. For example, chromVAR, Cicero, and Gene Scoring were designed to determine important marker genes, their regulatory logic, or to infer enriched TF binding sites within accessible chromatin regions. However, because clustering is a critical part of single-cell analysis and researchers frequently use output from all methods to produce clustering results [1], we felt that evaluating the clustering abilities using feature matrices produced by each method was a useful measure. An additional limitation of our study is that it is impossible to create a simulation framework that models an experimental outcome with perfect accuracy. Several assumptions were made to enable our simulation of the data; these assumptions are described in the methods section of this manuscript, where we detail explicitly how the simulated data was generated.

Interestingly, we learnt that some combinations of feature matrices with the simple clustering approaches included in our benchmarking framework perform even better than the original combination proposed by the respective authors. This highlights the value of this dual-characterization (*user* vs *designer perspective*) and provides a summary of both perspectives to the readers.

We believe it is important to stress the distinction between biological realities and computational performance, especially in the context of unsupervised clustering. A big and critical assumption (or hope) of our field is that an unsupervised clustering procedure will provide clustering solutions that recapitulate different populations corresponding to different cell types/states. Given that for several real datasets the ground truth is not known, a current compromise during the exploratory clustering analysis is to use known marker genes, sorted populations or known tissues to validate the clustering solutions based on classic metrics. If we embrace this assumption, keeping in mind that additional validation is required to truly delineate the subpopulation structure of a population of cells, the two views, biological and computational can be reconciled. Our benchmark procedure is aimed to provide some guidelines based on explorative analyses that are currently adopted in several published papers.

Looking forward, due to the wealth of data being produced by new scATAC technologies, we hypothesize that more powerful machine learning frameworks may be able to uncover complex *cis* and *trans* relationships that define cell-cell relatedness. Specifically, we anticipate autoencoder-like models that integrate genomic sequence context, gene body positions, and precise accessible chromatin information will yield information-rich features and that more advanced manifold learning methods will help to remove redundancy and better preserve heterogeneity within single cell populations. Such achievements may enable us to overcome the inherent sparsity and high dimensionality that characterizes scATAC-seq data.

### Conclusions

Our benchmarking results highlight SnapATAC, cisTopic, and Cusanovich2018 as the top performing scATAC-seq data analysis methods to perform clustering across all datasets and different metrics. Methods that preserve information at the peak-level (cisTopic, Cusanovich2018, Scasat) or bin-level (SnapATAC) generally outperform those that summarize accessible chromatin regions at the motif/k-mer level (chromVAR, BROCKMAN, SCRAT) or over the gene-body (Cicero, Gene Scoring). In addition, methods that implement a dimensionality reduction step (BROCKMAN, cisTopic, Cusanovich2018, Scasat, SnapATAC) generally show advantages over the other methods without this important step. SnapATAC is the most scalable method; it was the only method capable of processing more than 80,000 cells. Cusanovich2018 is the method that best balances analysis performance and running time.

Taken together, our manuscript provides a framework for evaluating and benchmarking new and existing methodologies as well as provides important guidelines for the analysis of scATAC-seq data. Importantly, we provide more than 100 well organized and documented Jupyter notebooks to illustrate and reproduce all the analyses performed in this benchmarking work. We believe our systematic analysis could guide the development of computational approaches aimed at solving the remaining challenges associated with analyzing scATAC-seq datasets.

## Methods

Our assessment of methods was based on public scATAC-seq datasets made available in public repositories by the respective authors (see **Data and code availability**). As such, we refer to the original publications for further details on experimental design and data pre-processing/alignment. For peak calling, we used the ENCODE pipeline (https://www.encodeproject.org/atac-seq/) except for the 10X PBMCs data for which peaks were already available through the Cell Ranger pipeline optimized for this technology. Whenever changes were required for running a given method, those are noted in the respective sections.

### Datasets

#### Human hematopoiesis I (Buenrostro et al. 2018)

This dataset comprised of 10 FACS-sorted cell populations from CD34^+^ human bone marrow, namely, hematopoietic stem cells (HSCs), multipotent progenitors (MPPs), lymphoid-primed multipotent progenitors (LMPPs), common-myeloid progenitors (CMPs), granulocyte-macrophage progenitors (GMPs), megakaryocyte-erythrocyte progenitors (MEPs), common-lymphoid progenitor (CLPs), plasmacytoid dendritic cells (pDCs), monocytes, and an uncharacterized CD34^+^ CD38^−^ CD45RA^+^ CD123^−^ cell population. A total of 2,034 cells from 6 human donors were used for analysis. A peak file (including 491,437 peaks) obtained from bulk ATAC-seq dataset was provided.

#### sci-ATAC-seq mouse tissues (Cusanovich et al. 2018)

This dataset comprises cells from 13 tissues of adult mouse, namely, bone marrow, cerebellum, heart, kidney, large intestine, liver, lung, prefrontal cortex, small intestine, spleen, testes, thymus, and whole brain, with over 2,000 cells per tissue. A total of 81,173 cells from 5 mice were used for analysis. A subset was obtained by randomly down-sampling 15% cells from each tissue and was comprised of 12,178 cells.

#### Human hematopoiesis II (10X PBMCs)

This dataset is composed of peripheral blood mononuclear cells (PBMCs) from one healthy donor. A total of 5,335 cells were used for analysis.

#### Simulated scATAC-seq datasets

In order to evaluate and benchmark various approaches, we generated synthetic (labeled) data from down-sampling 18 FACS-sorted bulk populations that were previously described [28]. For ease of interpretation, we considered only 6 isolated populations (HSC, CMP, NK, CD4, CD8, Erythroblast). For the erythropoiesis simulation, eight additional populations (P1-P8) originally described in [17] were also considered.

Our simulation framework starts with a peak x cell type counts matrix (from bulk ATAC-seq) and generates a single-cell counts matrix (*C*) for an arbitrary number of synthetic single cells. Explicitly, for a simulated single cell *j* and corresponding peak *i* from bulk cell type *t*, we seek to generate *c_i,j_* where *c_i,j_* ∈ Error! Bookmark not defined., noting that these values correspond to possible observations in a diploid genome. Next, we define the rate 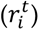 at which the peak *i* is prevalent in the bulk ATAC-seq data for cell type *t*. This rate is determined by the ratio of reads observed in peak *i* over the sum of all reads. Assuming a total of *k* peaks for the matrix *C* and for user-defined parameters *q* (noise parameter; *q* ∈ [0, 1]) and *n* (number of simulated fragments), we define *c_i,j_* as follows:

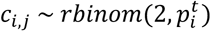

where

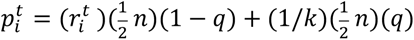

Intuitively, the parameter 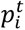 defines the probability that a count will be observed in peak *i* for a single cell. Additionally, 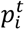 can be decomposed into the sum two terms. As *q* → 0, the first term dominates, and the probability of observing a count in peak *i* is simply the scaled probability of the ratio of reads for that peak from the bulk ATAC-seq data 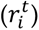. Thus, when *q* = 0, the simulated data has no noise. Conversely as *q* → 1, the second term dominates, and 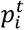 reduces to a flat probability that is no longer parameterized by the peak *i* or cell type *t* and thus represents a random distribution of *n* fragments into *k* peaks.

For bone marrow-based simulations we simulated 200 cells per labeled cell type while for erythropoiesis-based simulation we simulated 100 cells per labeled cell type. Eventually we have 1,200 cells for each simulated dataset. In the base simulations, we parametrized *n* = 2,500 fragments in peaks in expectation for all cells. For additional simulations that compared different data coverages, we set *n* to various values (5000, 2500, 1000, 500, 250 respectively) to benchmark this effect. To evaluate the effect of noise in our simulation, we set *q* to three values (0, 0.2, 0.4) to benchmark the robustness to noise. At values of *q* > 0.4, no method could reliably separate all the subpopulations. Finally, since our simulation started at the reads in peaks level, for some methods, the core algorithm associated with the method was extracted in order to benchmark it in this setting. Additionally, full code to reproduce these simulated dataset matrices has been made available with our online code resources.

#### Peak calling

For real datasets, peaks were called using the ENCODE ATAC-seq processing pipeline (https://www.encodeproject.org/atac-seq). Briefly, single-cells were aggregated into cell populations according to cell type, obtained either by FACS sorting or by tissue of origin. Peaks were called for each cell population and merged into a single file with bedtools [30].

#### Building the features matrix

##### BROCKMAN

This method starts by defining regions of interest, which will be scanned for *k*-mer content, as 50 bp windows around each transposon integration site and merging overlapping regions. Then, a frequency matrix of *k*-mers-by-cells is built by counting all possible gapped *k*-mers (for *k* from 1 to 8) within the previously defined windows. This frequency matrix is scaled so that each *k*-mer has mean 0 and standard deviation 1. Principal component analysis (PCA) is applied to the scaled *k*-mers-by-cells frequency matrix, and significant principal components (PCs) as estimated with the jackstraw method are selected to build a final features matrix for downstream analyses.

##### ChromVAR

This method starts by counting reads under chromatin-accessible peaks in order to build a count matrix of peaks-by-cells (X). Then, a set of chromatin features such as transcription factor (TF) motifs or *k*-mers are considered. Reads mapping to each peak that contains a given TF motif (or k-mer) are counted in order to build a count matrix of motifs-by-cells or k-mers-by-cells (M). Moreover, a raw accessibility deviation matrix of motifs (or k-mers)-by-cells (Y) is generated by calculating the difference between M and the expected number of fragments based on X. Then, background peak sets are created for each motif to remove technical confounders. Background motifs-by-cells raw accessibility deviations are then used to calculate a bias corrected deviation matrix and to compute a deviation z-score used for downstream analyses.

##### cisTopic

This method starts by building a peaks-by-cells binary matrix by checking if a peak region is accessible, i.e., at least one read falls within the peak region. Then, latent Dirichlet allocation (LDA) is performed on this binary matrix and two probability distributions are generated, a topics-by-cells probability matrix and a regions-by-topics probability matrix. The former is the final features matrix for downstream analyses.

##### Cicero

This method defines promoter peaks as the union of annotated TSS minus 500 base pairs and macs2 defined peaks around the TSS. It takes as input the peaks-by-cells binary matrix. It also requires either pseudo temporal ordering or coordinates in a low dimensional space (t-SNE) so that cells can be readily grouped. It then computes the co-accessibility scores between sites using Graphical Lasso. To get the gene activity scores, it selects sites that are proximal to gene TSS or distal sites linked to them and weight them by their co-accessibility. Then all the sites are summed and weighted according to their co-accessibility to produce a genes-by-cells feature matrix that is used in this benchmarking analysis.

##### Gene Scoring

This method first constructs a peaks-by-cells count matrix and defines regions of interest as the 50kb upstream and downstream of gene TSSs. Then it finds the overlap between ATAC-seq peaks and TSS regions and the peaks are weighted by a function of the distance to the linked genes. Finally, the peaks-by-cells count matrix is converted into genes-by-cells weighted count matrix by multiplying the weighted peaks by genes matrix. The genes-by-cells weighted count matrix is the final features matrix for downstream analyses.

##### Cusanovich2018

This method starts by binning the genome into fixed-size windows (by default, 5kbp), and building a binary matrix from evaluating whether any reads map to each bin. Bins that overlap ENCODE-defined blacklist regions are filtered out, and the top 20,000 most commonly used bins are retained. Then, the bins-by-cells binary matrix is normalized and rescaled using the term frequency-inverse document frequency (TF-IDF) transformation. Next, singular value decomposition (SVD) is performed to generate a PCs-by-cells LSI score matrix, which is used to group cells by hierarchical clustering into different clades. Within each clade, peak calling is performed on the aggregated scATAC-seq profiles, and identified peaks are combined into a new peaks-by-cells binary matrix. Finally, the new peaks-by-cells matrix is transformed with TF-IDF and SVD as before to get a matrix of PCs-by-cells, which is the final features matrix for downstream analyses.

##### scABC

This method starts by building a peaks-by-cells count matrix of read coverage within peak regions. Then, the weights of cells are calculated by a nonlinear transformation of the read coverage within the peaks background, defined as a 500 kb region around peaks. Since the weights will be used as part of weighted K-medoids clustering to define cell landmarks and further perform finer re-clustering instead of normalizing the peaks-by-cells matrix, the feature matrix in scABC is defined as the peaks-by-cells count matrix.

##### Scasat

This method first constructs a peaks-by-cells binary accessibility matrix by checking if at least one read overlaps with the peak region. Then Jaccard distance is computed based on the binary matrix to get a cells-by-cells dissimilarity matrix. Multidimensional scaling (MDS) is further performed to reduce the dimension and to generate the final feature matrix for downstream analysis.

##### SCRAT

This method starts by aggregating reads from each cell according to different features (such as TF motifs or region of interest of each gene), and then building a count matrix of features-by-cells. The features-by-cells count matrix is normalized by library and region size to get the final feature matrix for downstream analyses.

##### SnapATAC

This method starts by binning the genome into fixed-size windows (by default 5kb) and estimating read coverage for each bin to build a bins-by-cells binary count matrix. Bins that overlap ENCODE-defined blacklist regions are filtered out, as well as those with exceedingly high or low z-scored coverage. Then, the bins-by-cell matrix is transformed into a cells-by-cells Jaccard index similarity matrix, which is further transformed by normalization and regressing out coverage bias between cells. Finally, PCA is applied to the normalized similarity matrix, and the top PCs are used to build a PCs-by-cells matrix that is the final features matrix for downstream analyses.

### Clustering

For this study we used three commonly used clustering methods: k-means, hierarchical clustering (with default ward linkage) as implemented in the scikit-learn library [31] and Louvain clustering (a community-detection-based method) [32, 33] as implemented in Scanpy [34], For both hierarchical clustering and k-means, we set the number of clusters to the number of unique FACS-sorted labels or known tissues. In the 10X PBMCs scATAC-seq dataset, which lacks the FACS-sorted labels, we instead set the number of clusters to 8 since this is the expected number of populations based on previous studies [19]. For the Louvain algorithm, we set the size of local neighborhood to 15 for all the datasets. Since Louvain method requires ‘resolution’ instead of the number of clusters and different number of clusters will affect the clustering evaluation, to make the comparison fair, we use the binary search algorithm on the ‘resolution’ (ranging from 0.0 to 3.0) to find the same number of clusters as the other two clustering methods. If the precise number of clusters did not match the desired value, the ‘resolution’ value inducing the closest number of clusters to the desired value was used.

### Metrics for evaluating clustering results

To evaluate clustering solutions for datasets with a known ground truth (i.e. for each cell we have a label that indicated the cell type) we used three well-established metrics: Adjusted Rand Index (ARI), Mutual information and Homogeneity. Briefly, for the Adjusted Rand Index (ARI) first the Random Index (RI) is defined as a similarity measure between two clusters considering all pairs of samples assigned in the same or different clusters in the predicted and true clustering. Then, the raw RI score is adjusted for chance in the ARI score as described in the following formula:

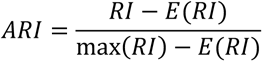

Where RI is the pre-computed random index and E is the expected random index.

Mutual Information is a measure of the mutual dependence between two variables. The Mutual Information value is computed according to the following formula, where *|Ui|* is the number of the samples in cluster *Ui* and *|Vj|* is the number of the samples in cluster *Vj:*

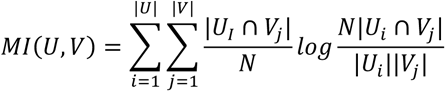

The homogeneity score is used to check if the algorithm used for the clustering can assign to each cluster only samples belonging to a single class. Its value *h* is bounded between 0 and 1, and a low value indicates low homogeneity and vice versa. The score is computed as follow:

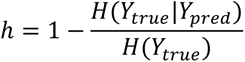

where *H(Ytrue|Ypred)* is the probability to assign true samples to a set of predicted samples, while *H(Ytrue)* are the labels of the samples.

To evaluate clustering solutions for the 10X PBMCs dataset we proposed a simple score called the Residual Average Gini Index (RAGI) and compared the accessibility of housekeeping genes with previously characterized marker genes [19]. We reasoned that a good clustering solution should contain clusters that are enriched for accessibility of different marker genes, and each marker gene should be highly accessible in only one or a few clusters. First, to quantify the accessibility of each gene in each cell we used the Gene Scoring approach described above. Briefly, the accessibility at each TSS is the distance-weighted sum of reads within or near the region. Second, to quantify the enrichment of each gene in each cluster of cells, we computed the mean of the accessibility values in all cells for each cluster. Third, based on the vector of mean accessibility values (one per cluster), we computed the Gini Index [35] for each marker gene. The Gini Index measures how imbalanced the accessibility of a gene is across clusters. This score is bound by [0, 1] where 1 means total imbalance (i.e. a gene is accessible in one cluster only) and 0 means no enrichment. This score has been previously used on scRNA-seq to perform clustering [36, 37]. As a control, we also calculated the Gini Index for a set of annotated housekeeping genes reported in (https://m.tau.ac.il/~elieis/HKG/HK_genes.txt). Housekeeping genes should show minimal specificity for any given cluster since, by definition, they are highly expressed in all cells. Based on the set of Gini Index values for marker and housekeeping genes we calculated several metrics: (i) the mean Gini Index for the two groups; (ii) the difference in means to assess the average residual specificity that a clustering solution has with respect to marker genes (this is our proposed RAGI metric); and (iii) the Kolmogorov Smirnov statistic and its p-value comparing the two groups of Gini Indices for marker and house-keeping genes. We sorted the methods based on the descending order of the differences in means (**Sup Table 13**); a positive value indicates that the marker genes on average separate the clusters better than uninformative housekeeping genes.

### Rare cell type-specific peak analysis

FACS-sorted bulk ATAC-seq data was downloaded and processed from a previously described resource [5]. For each simulation, we created a randomly-sampled set of 200 million unique (PCR-deduplicated) reads, which roughly represents a complexity similar to recommendations from the 10X Chromium scATAC-seq solution. Cell type-specific peaks were defined using the full dataset for each of the three cell types. Peaks were called using macs2 callpeak with custom parameters as in the ENCODE pipeline, i.e. *“--nomodel --shift - 100 --extsize 200*” to account for Tn5 insertions rather than read abundance when inferring peaks. Overlaps between the isolated minor population and the synthetic mixtures were computed using GenomicRanges[38] findOverlaps function, which is equivalent to bedtools[30] overlap. For each minor population (B-cell, CD4+ T-cell, Monocyte) and each prevalence (1, 5, 10, 20, 30%), each simulation was repeated 5 times for a total of 75 simulations. Reads from the other two (major) populations were sampled equivalently to make up the synthetic mixture for comparison.

### Data and code availability

All the results presented in this manuscript can be reproduced using the Jupyter notebooks available both at https://github.com/pinellolab/scATAC-benchmarking/ and in the supplementary material (**Sup Data**). For the analyzed real datasets, the *Buentrostro2018* dataset was downloaded from GEO accession GSE96772, the 10X PBMCs dataset was downloaded from https://support.10xgenomics.com/single-cell-atac/datasets/1.0.1/atac_v1_pbmc_5k, and the sci-ATAC-seq mouse dataset was downloaded from http://krishna.gs.washington.edu/content/members/ajh24/mouse_atlas_data_release/bams. For the simulated bone marrow dataset, data for the FACS-sorted bulk ATAC-seq populations were downloaded from GEO accession GSE119453. For the simulated erythropoiesis dataset, the additional populations were downloaded from GEO accession GSE115672.

## Author Contributions

H.C. and L.P. conceived this project and designed the framework with input from all the authors. H.C. and T.A. preprocessed data. H.C., C.L., T.A., M.E.V., S.P.G. and K.C. implemented scATAC-seq methods. H.C. and K.C. performed clustering analysis. H.C., T.A., L.P. performed clustering validation. C.L. simulated data. H.C. and C.L. analyzed simulated data. L.P. and J.D.B. provided guidance. All the authors wrote the manuscript.

## Acknowledgments

This project has been made possible in part by grant number 2018-182734 to L.P. from the Chan Zuckerberg Initiative DAF, an advised fund of Silicon Valley Community Foundation. L.P. is also partially supported by a National Human Genome Research Institute (NHGRI) Career Development Award (R00HG008399).

## Supplementary Notes

### Supplementary Note 1: Analysis of the simulated datasets

For all the synthetic datasets, the input is a peaks-by-cells raw count matrix generated as described in the Methods section. For all methods, we first order peaks based on the number of cells in which the peak is observed and select the top 8,000 peaks (making sure each of these peaks appear at least in one cell).

For BROCKMAN, we scanned for gapped k-mers (the default setting is used, i.e. length 1–8, all possible gaps) within peaks to calculate the scaled k-mer frequencies for each cell. For chromVAR, we used both TF binding motifs from the JASPAR database (human) or short k-mers (k=6) within peaks to score the accessibility deviation across cells. For Cicero, we run it with the default parameters to calculate gene activity scores. For cisTopic, we run it with the same parameters shown in their online tutorial (https://rawcdn.githack.com/aertslab/cisTopic/f628c6f60918511ba0fa4a85366ebf52db5940f7/vignettes/CompleteAnalysis.html). For *Cusanovich2018* we first binarize the count matrix and then perform the proposed TF-IDF transformation and SVD. For Gene Scoring, we select peaks overlapping with the regions of 50,000 bp upstream and downstream of TSSs as described in [1]. For scABC, since its feature matrix is the same as input matrix of peaks-by-cells, we instead run the steps of calculating the weights of cells that are used later for their proposed clustering approach. For Scasat, we first binarize the count matrix and then calculate Jaccard distance, followed by Multi Dimensional Scaling (MDS) with 10 dimensions (the same number of components as used for the Control-Naive). For SCRAT, the accessibility of TF binding motifs is summarized within peaks. We attempted to adjust for the library size and peak region length as suggested in the original study, however we noticed that this step dramatically penalizes this method performance in all the tested conditions (**Sup Fig. 1**). This step was therefore disabled for all the analyses performed with SCRAT. For SnapATAC, we use the fixed-size peaks as its bins. The Jaccard Index is normalized with the authors’ proposed method, *normOVE*. For methods that implement PCA step, we use the elbow plot to decide the optimal number of PCs. For methods that do not implement a step of dimensionality reduction, we use the R package *irlba* [2] to perform PCA.

All the notebooks detailing the exact procedures are available at https://github.com/pinellolab/scATAC-benchmarking/tree/master/Synthetic_Data.

### Supplementary Note 2: Analysis of the Buenrostro2018 dataset

For this dataset we started with aligned files in bam format (one per cell). We removed duplicated reads using the function *MarkDuplicates* version 2.20.2 with the option *REMOVE DUPLICATES = TRUE* from Picard (https://broadinstitute.github.io/picard/).

For the methods that do not provide an explicit function to read in bam files and count reads under peaks, including Cicero, Cusanovich2018, GeneScoring, Scasat, and Control-Naïve, we used a simple script to obtain a common peaks-by-cells raw count matrix (e.g. https://github.com/pinellolab/scATAC-benchmarking/tree/master/Real_Data/Buenrostro_2018/run_methods/Cusanovich2018/count_reads_peaks.sh). For the methods that implement the same strategy to filter peaks based on their frequency, including Cicero, Control-Naive, Cusanovich2018, GeneSoring, Scasat, and scABC, we filter out peaks that are observed in less than 1% of cells. For chromVAR, we run its function *filterPeaks* with the default setting to filter out peaks based on the minimum number of fragments and merge overlapping peaks. For the methods that implement a PCA step, including BROCKMAN, Control-Naïve, Cusanovich2018, and SnapATAC, we decided the number of PCs based on the elbow plot. For Scasat, which implements MDS, we set the number of dimension as 15 according to its tutorial https://github.com/ManchesterBioinference/Scasat/blob/master/ScAsAT_functions_Buenrostro_All_Bam_Together.ipynb. For cisTopic, the number of topics (dimensions) is decided by its function *selectModel* with default settings.

For the clustering analysis, we set the expected number of clusters as the number of FACS-sorting labels (10 in this case). For k-means, we use the *k-means++* to select the initial cluster centers. For hierarchical clustering, we use the *Ward* linkage based on Euclidean distance. Both k-means and hierarchical clustering are implemented in scikit-learn package[3]. For Louvain, we set the number of neighbors to 15 and the resolution is decided using a binary search with 20 steps that explores values of the resolution parameter in the interval 0~3. The Louvain algorithm used is implemented in Scanpy[4]. For the UMAP visualization, we run the function ‘umap’ from the R package *umap* with default settings.

All the notebooks for this analysis are available at https://github.com/pinellolab/scATAC-benchmarking/tree/master/Real_Data/Buenrostro_2018 and https://github.com/pinellolab/scATAC-benchmarking/tree/master/Real_Data/Buenrostro_2018_bulkpeaks.

### Supplementary Note 3: Analysis of 10x PBMCs dataset

For this dataset, we started with a single merged bam file downloaded from the 10x website and preprocessed with Cell Ranger: https://support.10xgenomics.com/single-cell-atac/datasets/1.0.1/atac_v1_pbmc_5k. We noticed that all the methods except SnapATAC don’t support this format i.e. a single bam file for multiple cells. Therefore, using the cell barcodes passing quality filtering from Cell Ranger, we split this file in multiple bam files, one per cell recovering 5,335 single-cell bam files. We also removed duplicate reads using Picard and performed UMAP visualization as discussed in **Supplementary Note 2**. For the clustering analysis, we set the expected number of clusters as the number of putative cell types (8 in this case) as previous studies suggested [5, 6].

All the notebooks are available at https://github.com/pinellolab/scATAC-benchmarking/tree/master/Real_Data/10x_PBMC_5k.

### Supplementary Note 4: Analysis of the sci-ATAC-seq mouse dataset

For this dataset, we started with multiple merged bam file from 17 samples across 13 tissues downloaded from http://krishna.gs.washington.edu/content/members/ajh24/mouse_atlas_data_release/bams. For each tissue we performed the same steps as in 10x PBMCs dataset to decompose the single merged bam file to multiple single cell bam files (81,173 bam files). The downloaded bam files were already deduplicated. The downsampled dataset of 12,178 cells is generated by randomly selecting 15% from each sample.

The scATAC-seq methods and UMAP visualization are implemented as in **Supplementary Note 2**. For the clustering analysis, we set the expected number of clusters as the number of tissues (13 in this case).

All the notebooks are available at https://github.com/pinellolab/scATAC-benchmarking/tree/master/Real_Data/Cusanovich_2018 and https://github.com/pinellolab/scATAC-benchmarking/tree/master/Real_Data/Cusanovich_2018_subset.

### Supplementary Note 5: Memory requirements and implementation choices

As mentioned in the main text, SnapATAC is the only methods that allows to process successfully large datasets, as the sciATAC-seq mouse dataset with ~80000 cells. Here we investigate why the other methods failed to analyze this large dataset. We hypothesize that main reason is related to the way the methods load/process the data in memory. In fact, we discovered that several methods require to load the entire dataset in the central memory (RAM).

BROCKMAN, Cicero and Gene Scoring try to load the entire dataset in memory using the *read.table* function or the *fread* function within the *data.table* package in R. Other methods such as: Cusanovic, Scrat, chromVAR, scABC and Scasat, store the entire dataset in memory within a *Matrix* object in R. CisTopic, has an optimized step to map the reads into the genome using the *Rsubread* function. This function creates a hash table of the entire genome and allows the user to select the amount of memory to use. At the end, the entire dataset is stored in the computer memory in a *CisTopicObject* data structure.

SnapATAC, preprocess the entire dataset and store it a *.snap* file. This file is based on the HDF5 technology that allows out of core computation. In SnapATAC the Python library h5py (a wrapper for HDF5 core library) is used to create the custom snap file format. More information about this custom file are available here: https://github.com/r3fang/SnapTools/blob/master/docs/snap_format.docx.

### Supplementary Note 6: End-to-end user-perspective clustering analysis

For the methods that explicitly implement the step of clustering in their tutorials, including Cusanovich2018, cisTopic, SnapATAC, scABC, Cicero, and Scasat, in addition to the three clustering methods used in this benchmark framework, we also performed the clustering analysis as shown in each tutorial. For Cusanovich2018, we followed the tutorial at http://atlas.gs.washington.edu/fly-atac/docs/ and used density peak algorithm [7] to identify clusters. For cisTopic, we followed the tutorial at https://rawcdn.githack.com/aertslab/cisTopic/f628c6f60918511ba0fa4a85366ebf52db5940f7/vignettes/CompleteAnalysis.html and used ward hierarchical clustering to cluster cells. For SnapATAC, we followed tutorial at https://github.com/r3fang/SnapATAC/blob/master/examples/10X_P50/README.md and used Leiden algorithm to cluster cells. For scABC we followed the tutorial at https://github.com/SUwonglab/scABC/blob/master/vignettes/ExampleWorkflow.html and used weighted k-medoids clustering. For Cicero we followed the tutorial at https://www.bioconductor.org/packages/devel/bioc/vignettes/cicero/inst/doc/website.html. To be consistent with the feature matrix used in the benchmarking framework, instead of using its default peaks-by-cells count matrix, we used gene activity scores as the input of clustering analysis. After reducing the dimensionality with tSNE, density peak clustering algorithm is used to cluster cells. For Scasat, we follow the tutorial at https://github.com/ManchesterBioinference/Scasat/blob/master/ScAsAT_functions_Buenrostro_All_Bam_Together.ipynb and use *ward.D2* hierarchical clustering for clustering.

We run all the six methods on three real datasets, Buenrostro2018, 10x PBMCs 10x dataset, sci-ATAC-seq mouse dataset. For Buenrostro2018 and sci-ATAC-seq mouse dataset, we specified the number of clusters as the number of FACS-sorted labels and the number of tissues respectively. For 10x PBMCs, we specified the number of clusters as 8 as suggested by the previous studies [5, 6].

### Supplementary Note 7: Running time

For the real datasets, we recorded the execution time of each method to generate a feature matrix starting from an aligned and deduplicated bam file. We noticed that not all the methods provide specific functions to read in bam files. Some methods only start with features by cells raw matrix (e.g. Cicero). In addition, the functions to count reads of some methods were not generalizable across the different scATAC-seq techniques (e.g. *Cusanovich2018*). Therefore, to make a fair comparison we used a common script (https://github.com/pinellolab/scATAC-benchmarking/blob/master/Real_Data/Buenrostro_2018/run_methods/Control/count_reads_peaks.sh) to obtain the peaks by cells matrix starting from bam files for the following 4 methods: Control-Naïve, *Cusanovich2018*, Gene Scoring, Scasat. BROCKMAN, perform two steps to obtain the final feature matrix (q bash script to count k-mer frequency and a R function to assemble the matrix), so we are considering the sum of their running times. Similarly, the running time for SnapATAC is based on two steps: the *snaptools* utility that converts a bam to the required *.snap* format and the R function that generates the feature matrix.

For the simulated datasets, we recorded the execution time of generating feature matrices starting from a simulated peaks-by-cell count matrix. For scABC, since its feature matrix is the same as the input, to have a useful running time, we instead record the time to calculate the cells weights, which are necessary for downstream analysis.

We also assessed the scalability of the methods with respect to the read coverage (250, 500, 1000, 2500 and 5000 fragments per peaks). We observed that the running time of most methods is not affected by the read coverage. This is not surprising given that our simulation them number of peaks is fixed, so the dimensionality of the matrix is unchanged. However, for cisTopic, we noticed an exponential increase in running times as we increase the number of fragments (**Sup Fig. 22**). We assume this might be due to the topic modelling approach used by cisTopic since it tries to learn the probability distribution over the regions for each topic while high coverage will result in the increase in the number of accessible regions.

## Supplementary Figures

**Figure S1.**
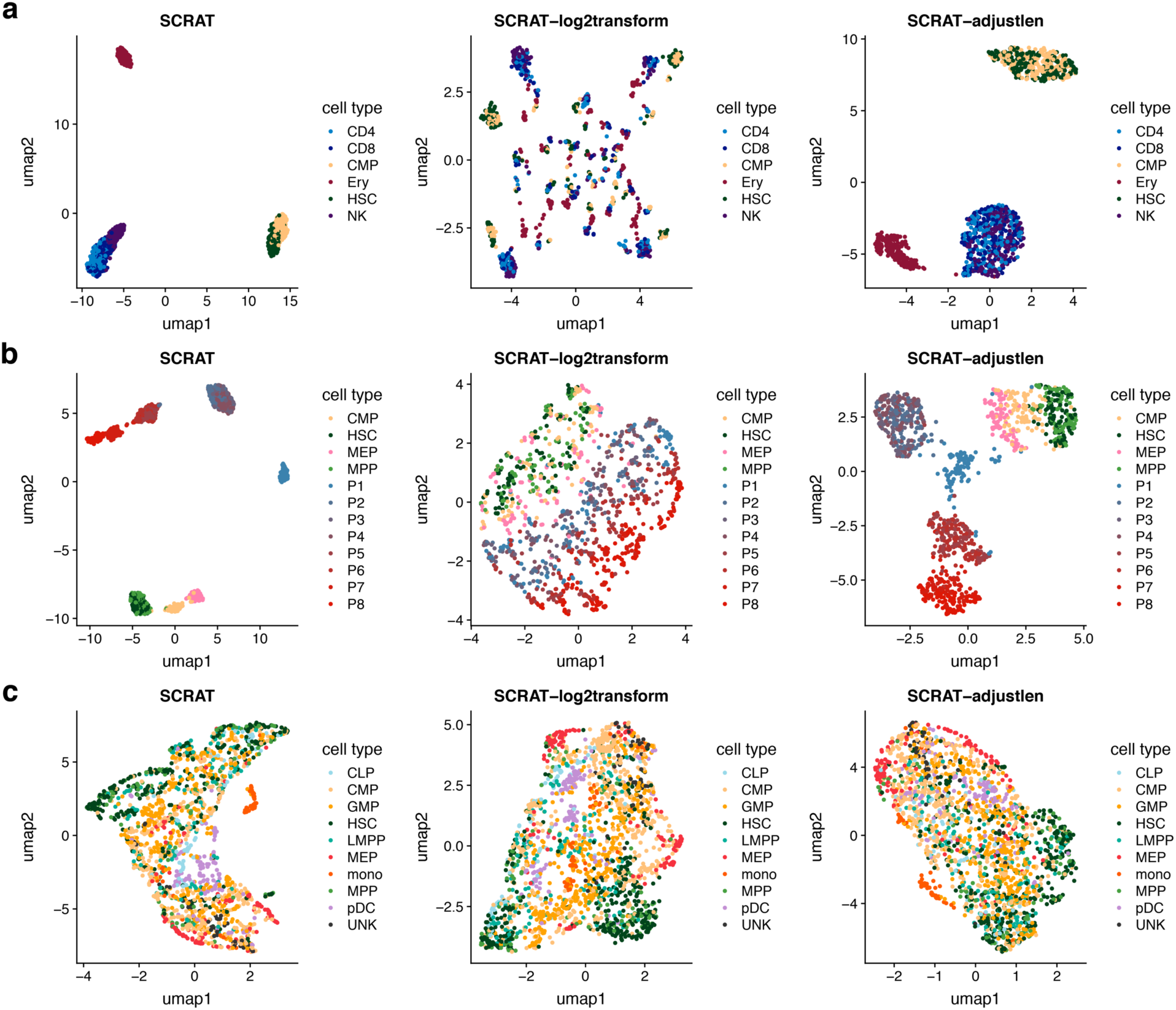
UMAP visualization of cells based on SCRAT feature matrix with different parameter settings (Left: log2transform=FALSE, adjustlen=FALSE. Middle: log2transform=TRUE, adjustlen=FALSE. Right: log2transform=FALSE, adjustlen=TRUE) in three datasets. **(a)** simulated bone marrow dataset at a noise level of 0.2 with a coverage of 2,500 fragments **(b)** simulated erythropoiesis dataset at a noise level of 0.2 with a coverage of 2,500 fragments **(c)** *Buenrostro 2018* dataset.

**Figure S2.**
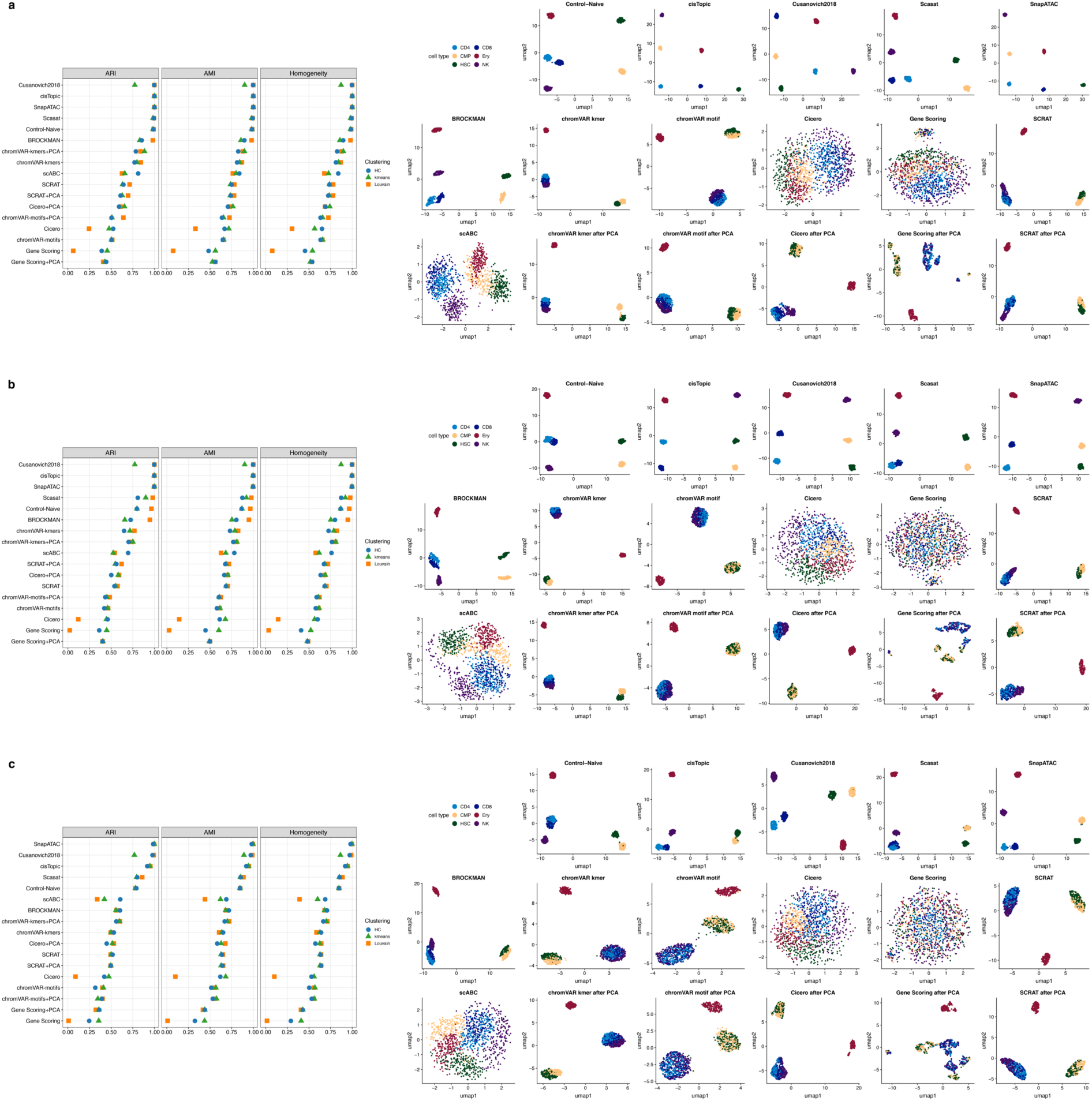
Clustering evaluation according to AMI, ARI and Homogeneity metrics (***left***) and UMAP visualization of cells colored by known cell labels (***right***) in simulated bone morrow datasets with a coverage of 2,500 fragments at **(a)** no noise (0), **(b)** moderate noise (0.2) and **(c)** high noise (0.4).

**Figure S3.**
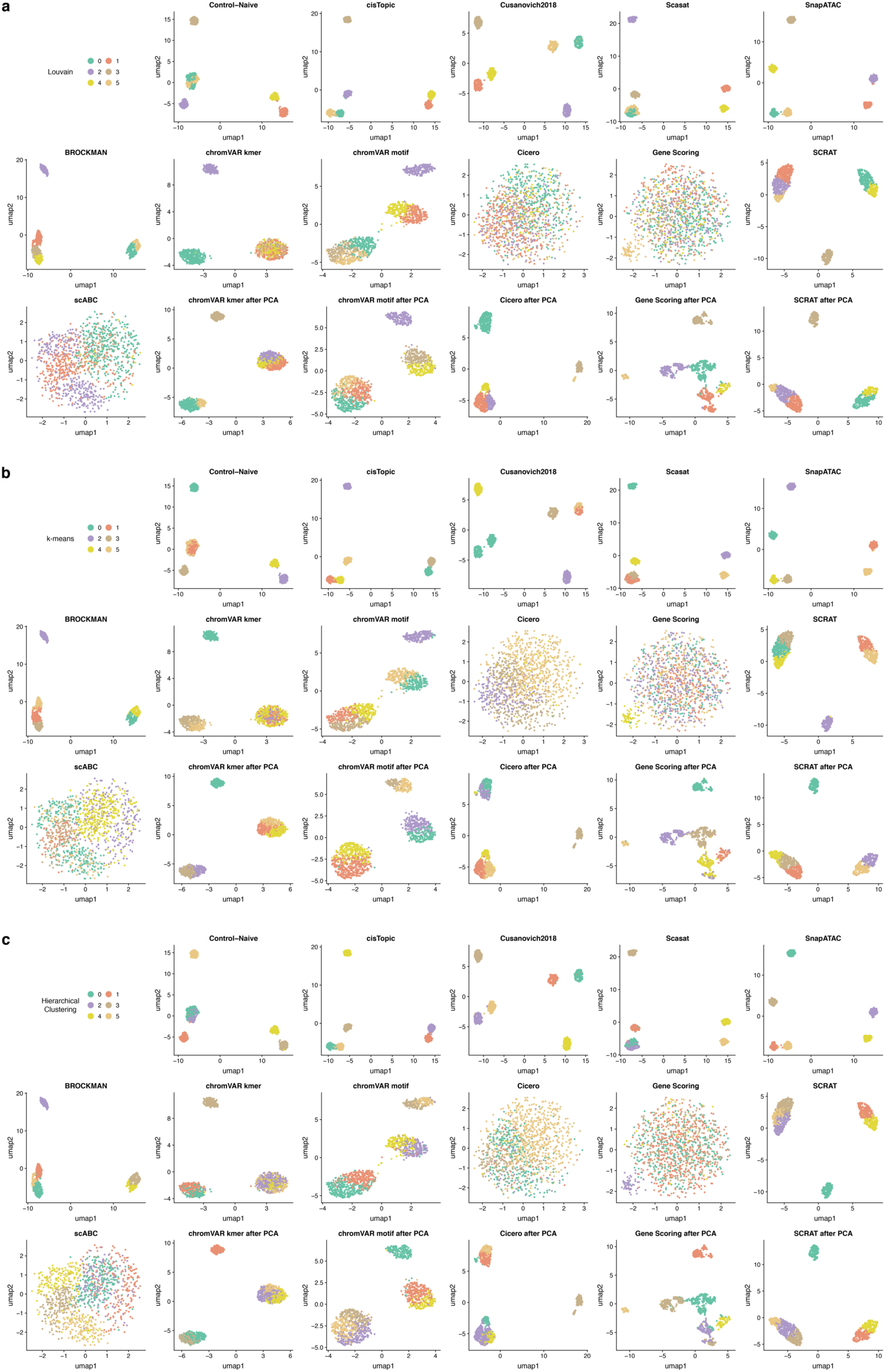
UMAP visualization of cells colored by clustering solution on the simulated bone marrow dataset with a noise level of 0.4 and a coverage of 2,500 fragments using **(a)** Louvain algorithm, **(b)** k-means clustering, and **(c)** hierarchical clustering (HC).

**Figure S4.**
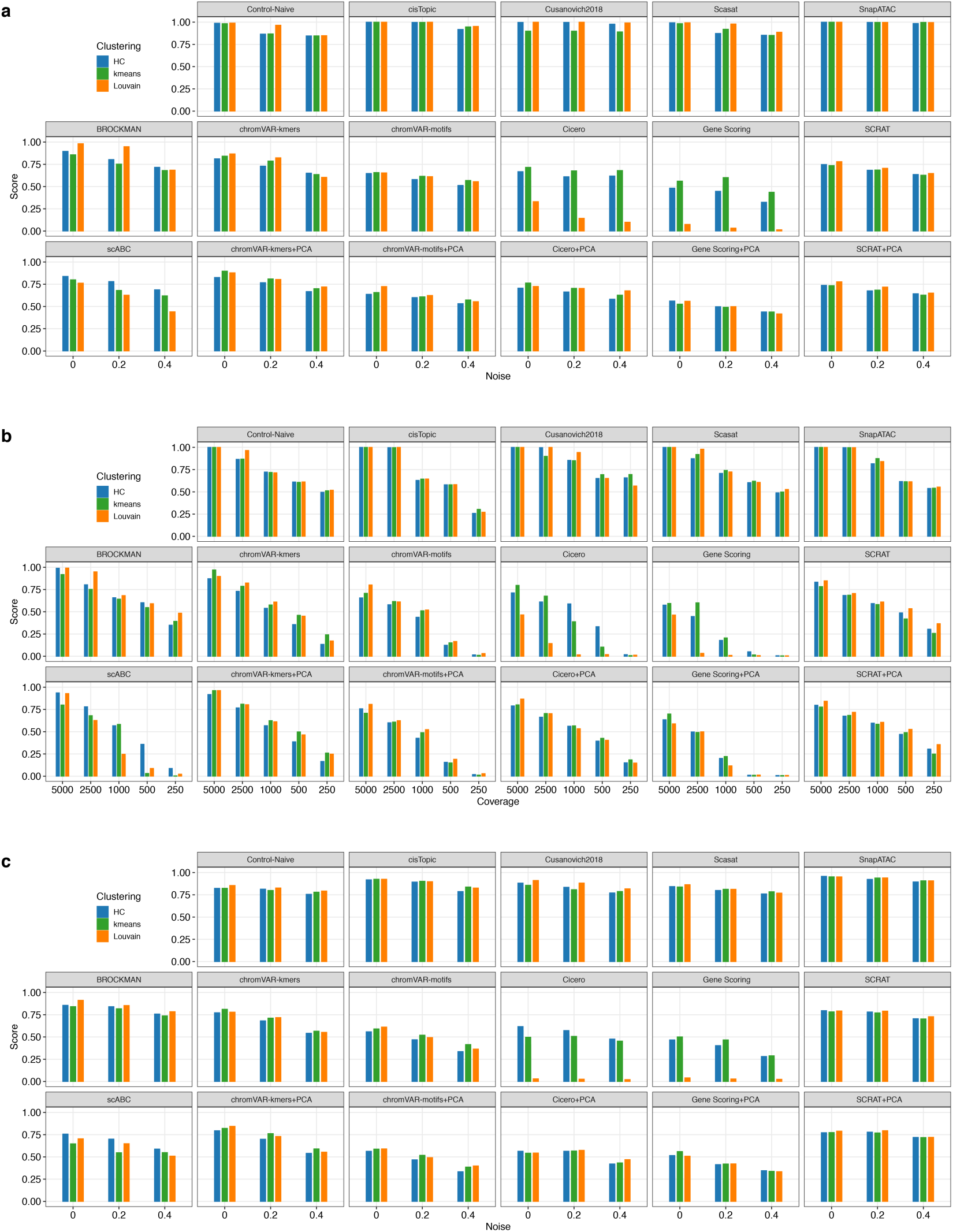
Summary of clustering scores at different noise levels and coverages based on three different clustering methods including hierarchical clustering (HC), k-means clustering and the Louvain algorithm. **(a)** clustering scores at noise levels of 0, 0.2, and 0.4 for the simulated bone marrow dataset with a coverage of 2,500. **(b)** clustering scores at the coverages of 5000, 2500, 1000, 500, 250 in the simulated bone marrow dataset at the noise level of 0.2. **(c)** clustering scores at the noise levels of 0, 0.2, and 0.4 for the simulated erythropoiesis dataset with a coverage of 2,500.

**Figure S5.**
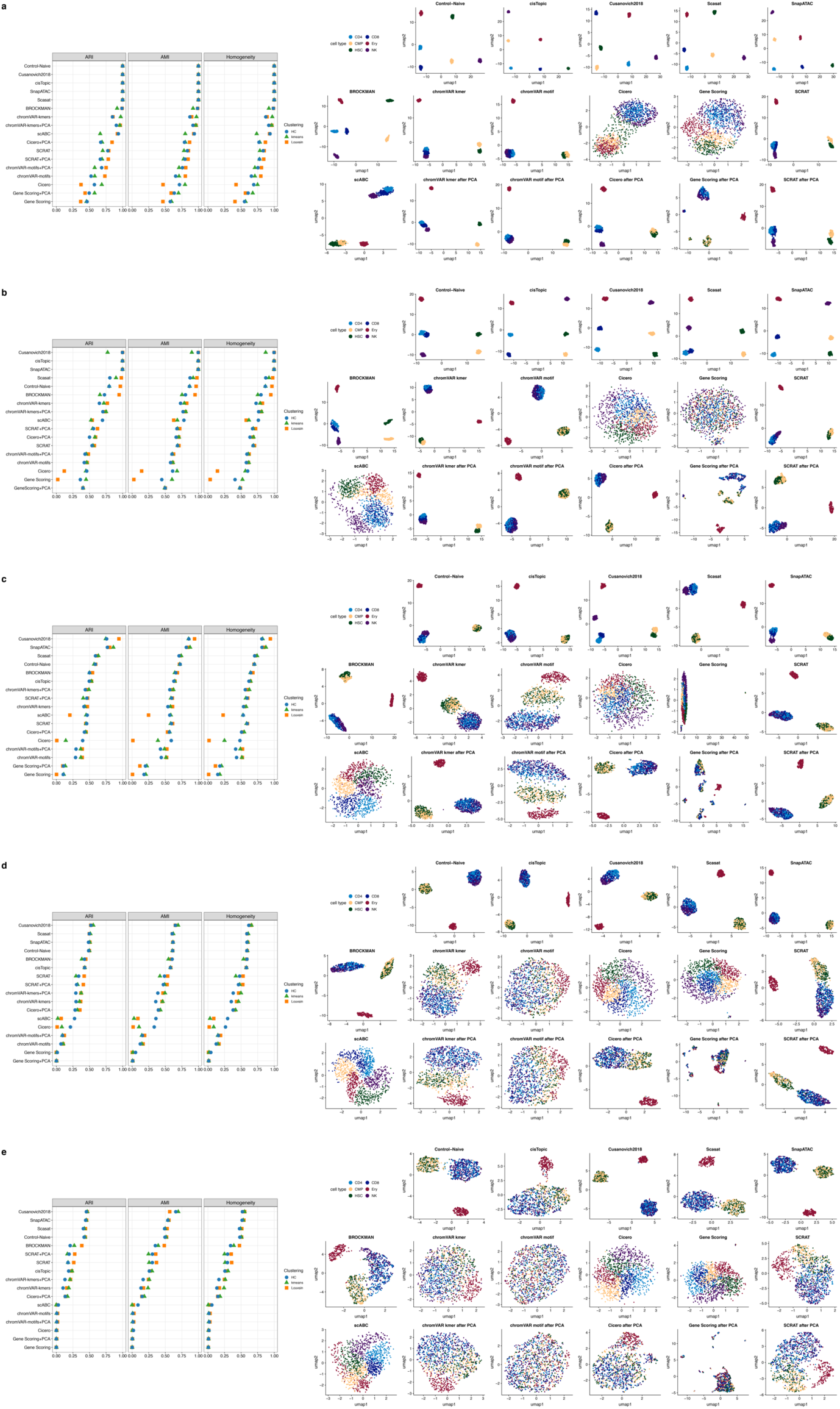
Clustering evaluation according to AMI, ARI and Homogeneity metrics (***left***) and UMAP visualization of cells colored by known cell labels (***right***) for the simulated bone marrow dataset with a noise level of 0.2 and varying coverages: **(a)** 5000 reads, **(b)** 2500 reads, **(c)** 1000 reads, **(d)** 500 reads, and **(e)** 250 reads.

**Figure S6.**
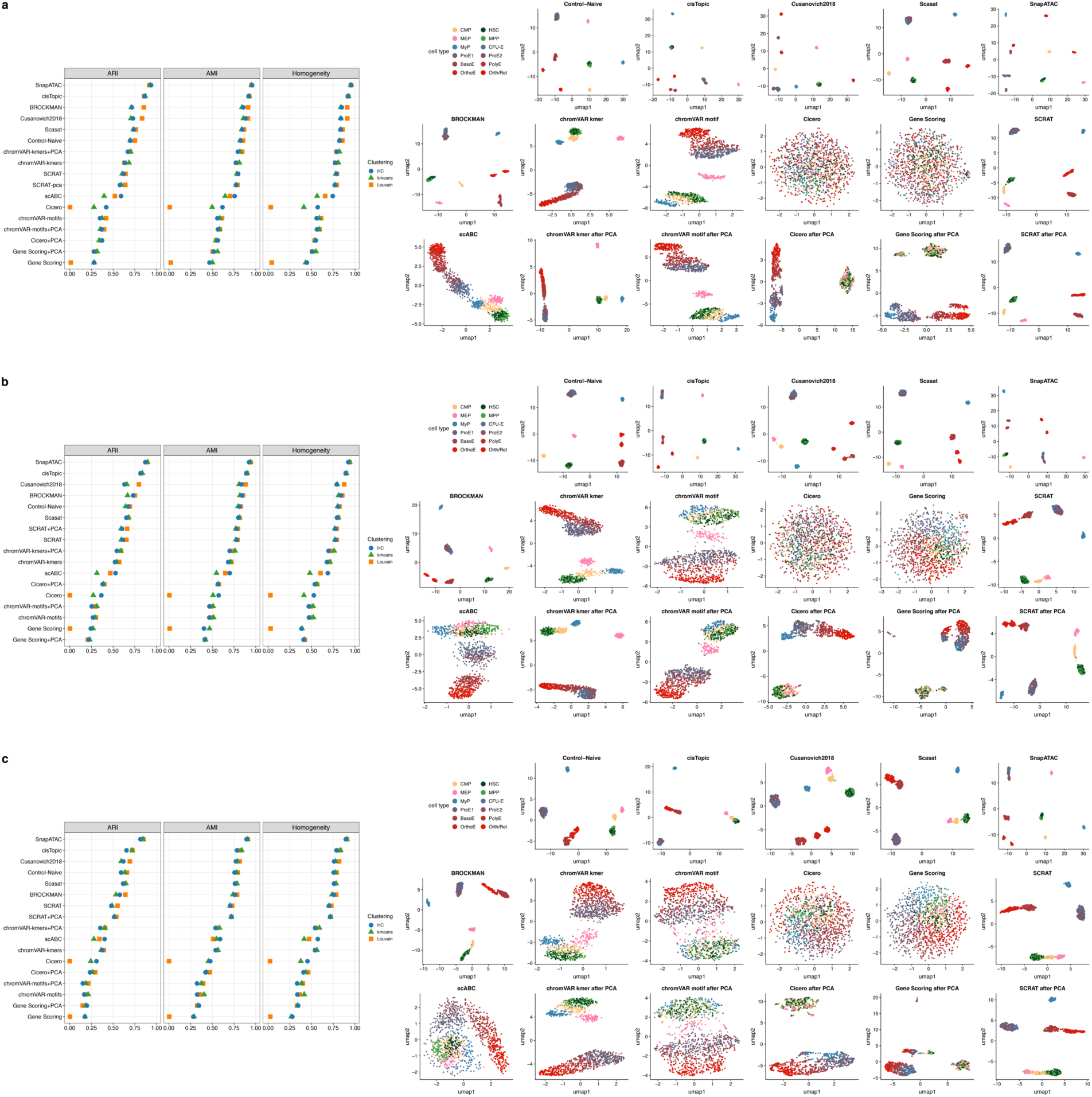
Clustering evaluation according to AMI, ARI and Homogeneity metrics (***left***) and UMAP visualization of cells colored by known cell labels (***right***) for the simulated erythropoiesis datasets with a coverage of 2,500 fragments and **(a)** no noise (0), **(b)** moderate noise (0.2) or **(c)** high noise (0.4).

**Figure S7.**
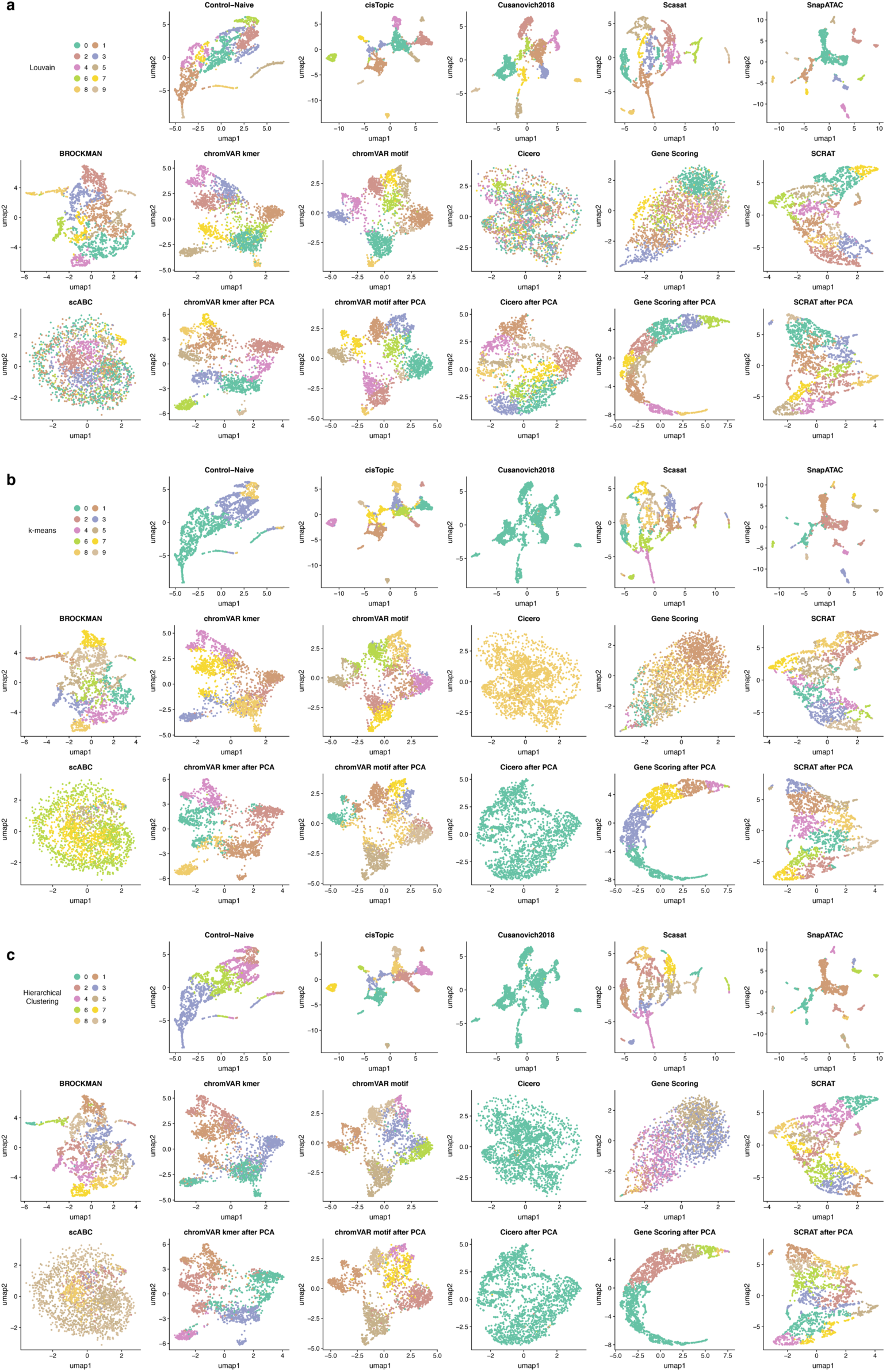
UMAP visualization of cells colored by the clustering solution on the *Buenrostro2018* dataset using **(a)** the Louvain algorithm, **(b)** k-means clustering, and **(c)** hierarchical clustering (HC).

**Figure S8.**
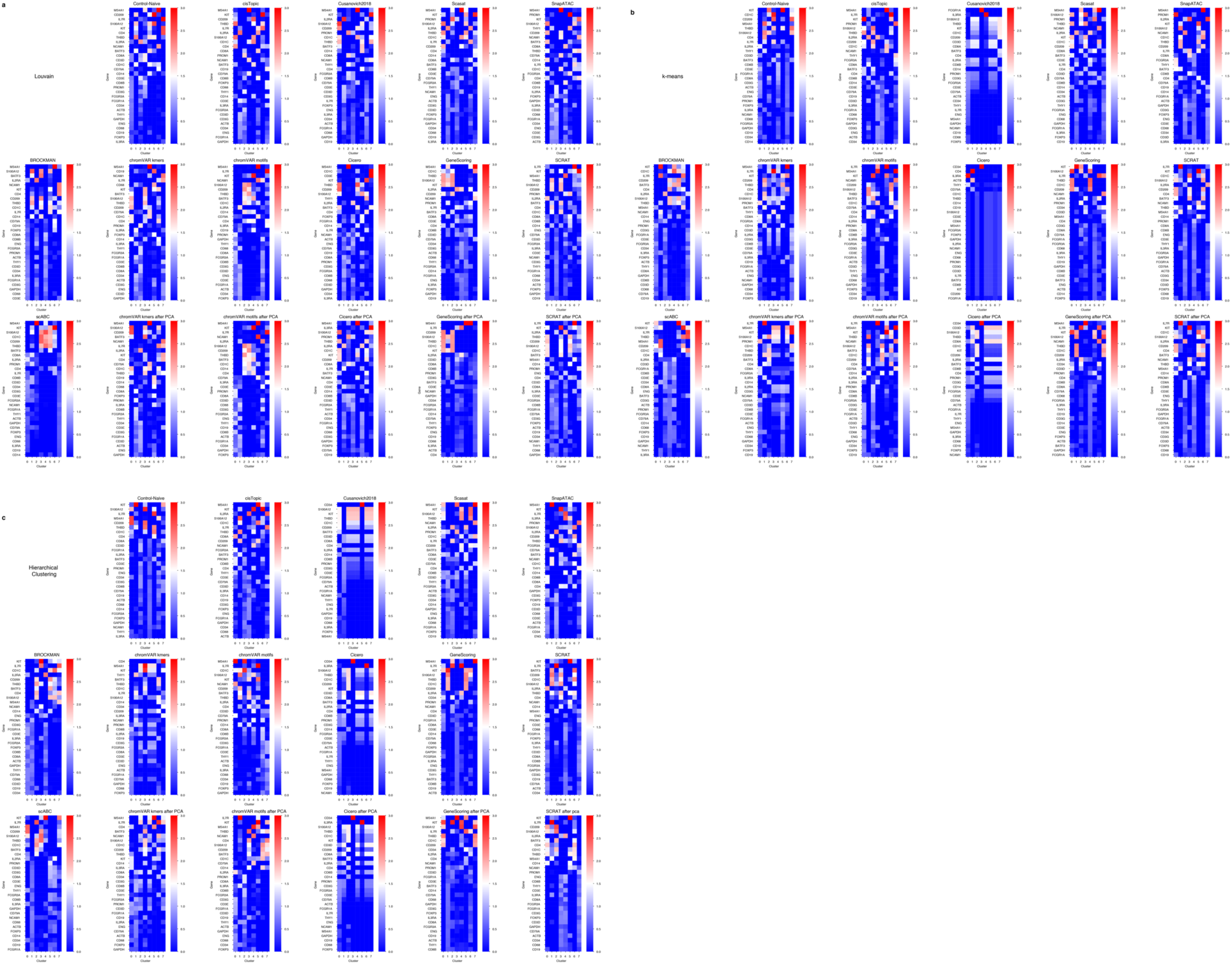
Heatmap for the average accessibility across clusters (columns) and the marker genes (rows) that are used to calculate the RAGI metric on the 10X PBMCs dataset. **(a)** Louvain clustering solution **(b)** k-means clustering solution **(c)** hierarchical clustering (HC) clustering solution.

**Figure S9.**
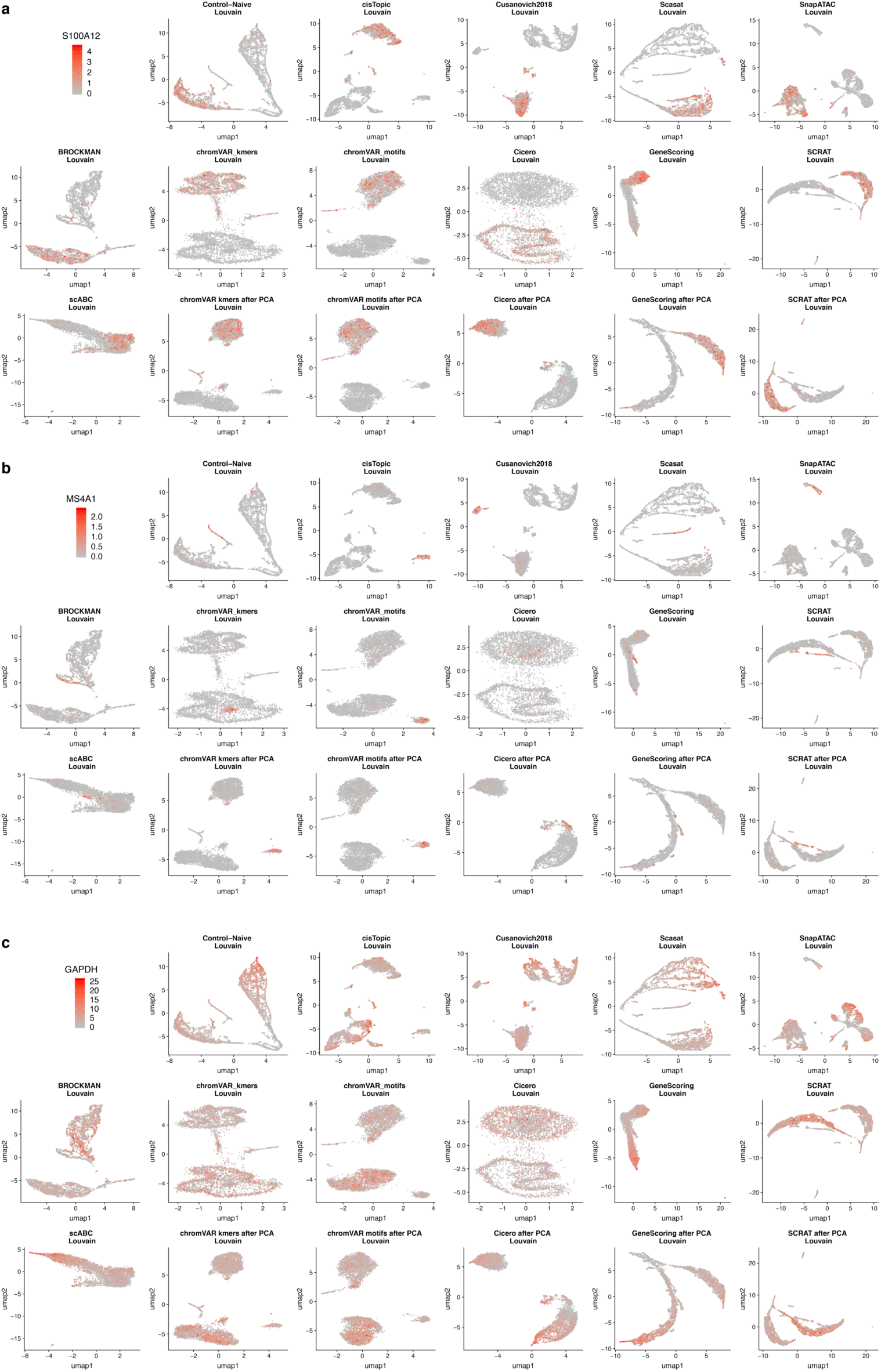
UMAP visualization of cells colored by the accessibility of marker genes: **(a)** S100A12 and **(b)** MS4A1 and **(c)** GAPDH (housekeeping gene) and on the 10X PBMCs dataset.

**Figure S10.**
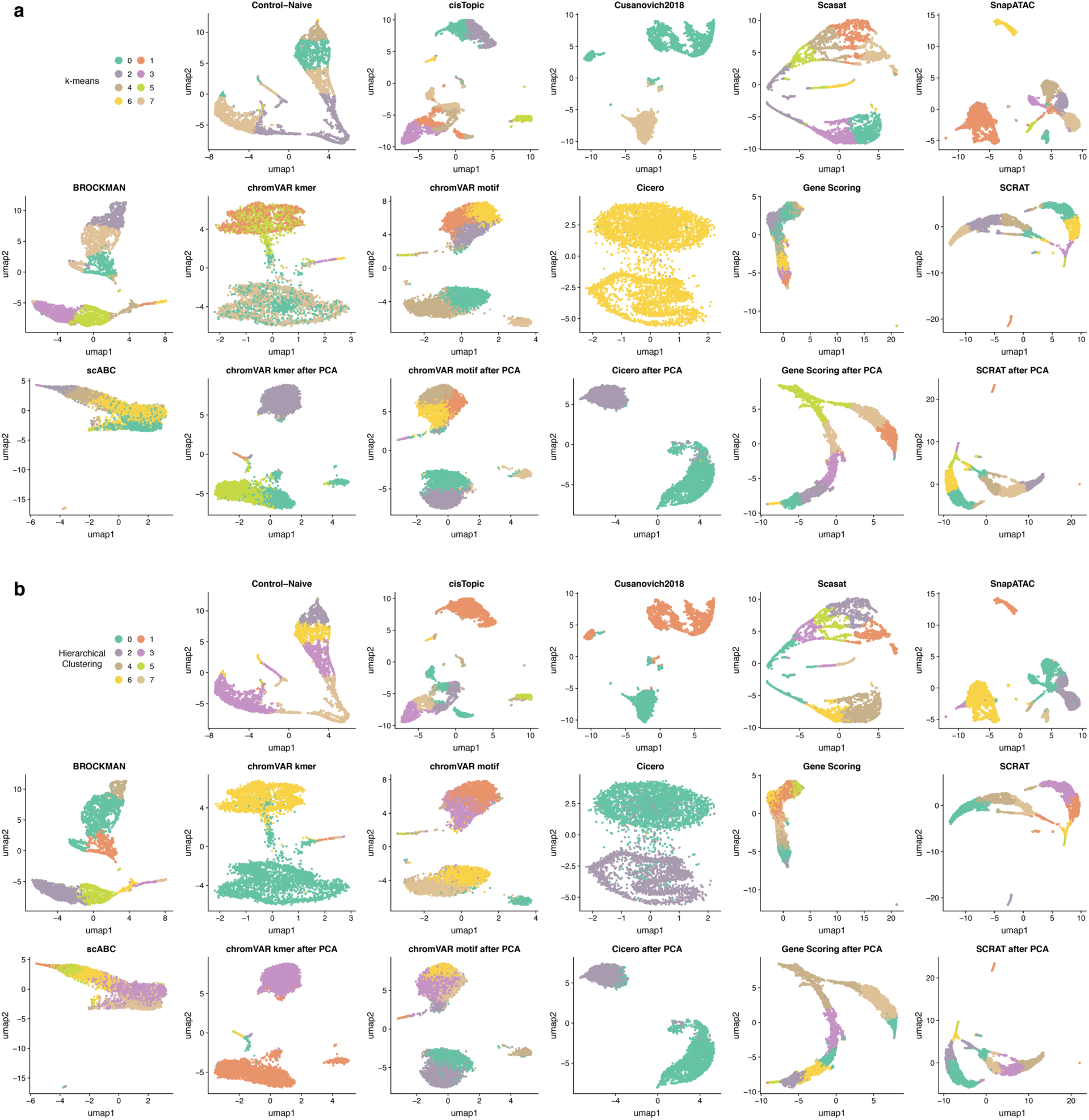
UMAP visualization of cells colored by the clustering solution on 10X PBMCs dataset using **(a)** k-means clustering and **(b)** hierarchical clustering (HC).

**Figure S11.**
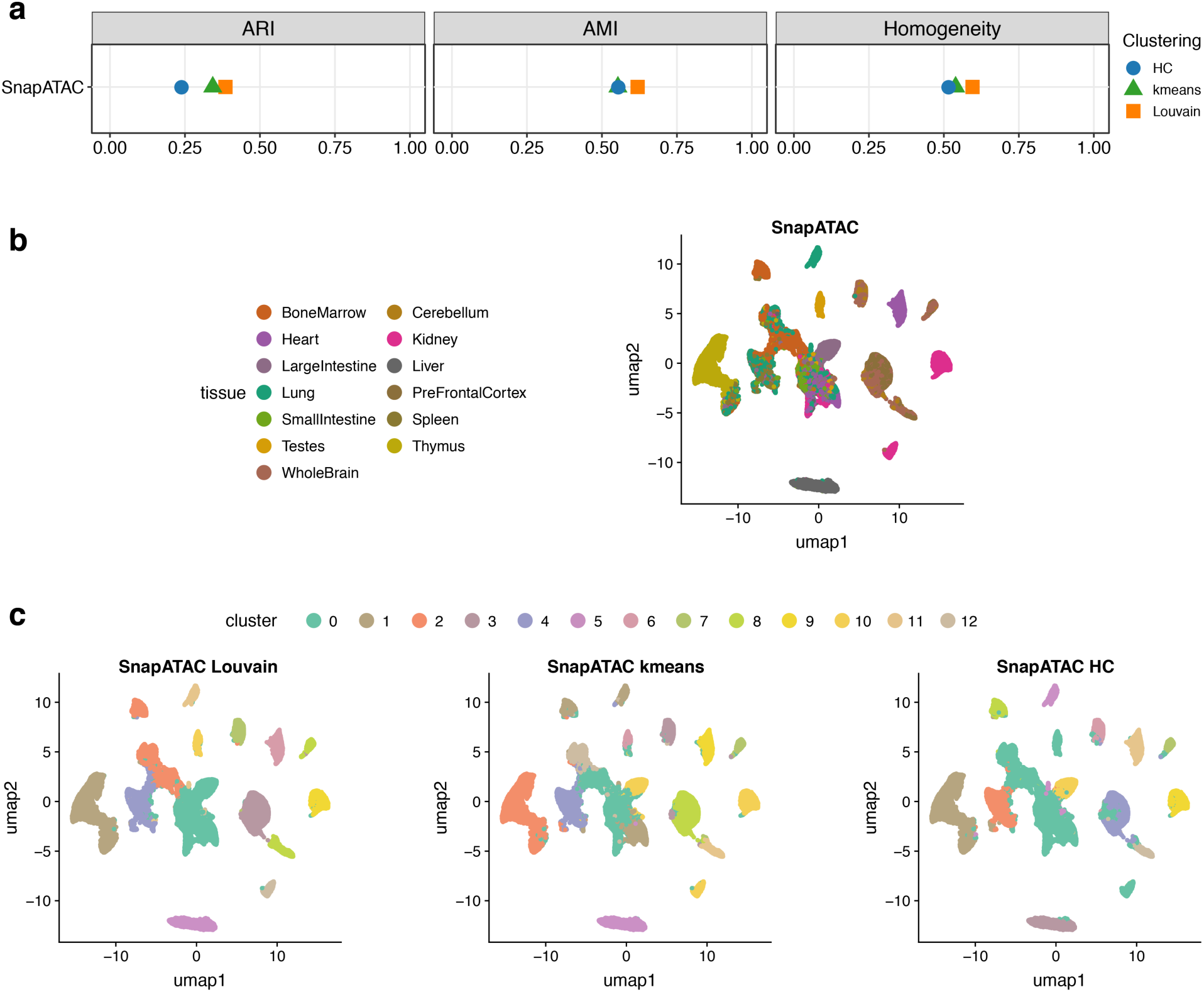
Assessment of SnapATAC on the full sci-ATAC-seq mouse dataset. **(a)** Clustering scores according to AMI, ARI and Homogeneity metrics **(b)** UMAP visualization of cells colored by the known tissues. **(c)** UMAP visualization of cells colored by three clustering solutions: the Louvain algorithm, k-means clustering, and hierarchical clustering (HC).

**Figure S12.**
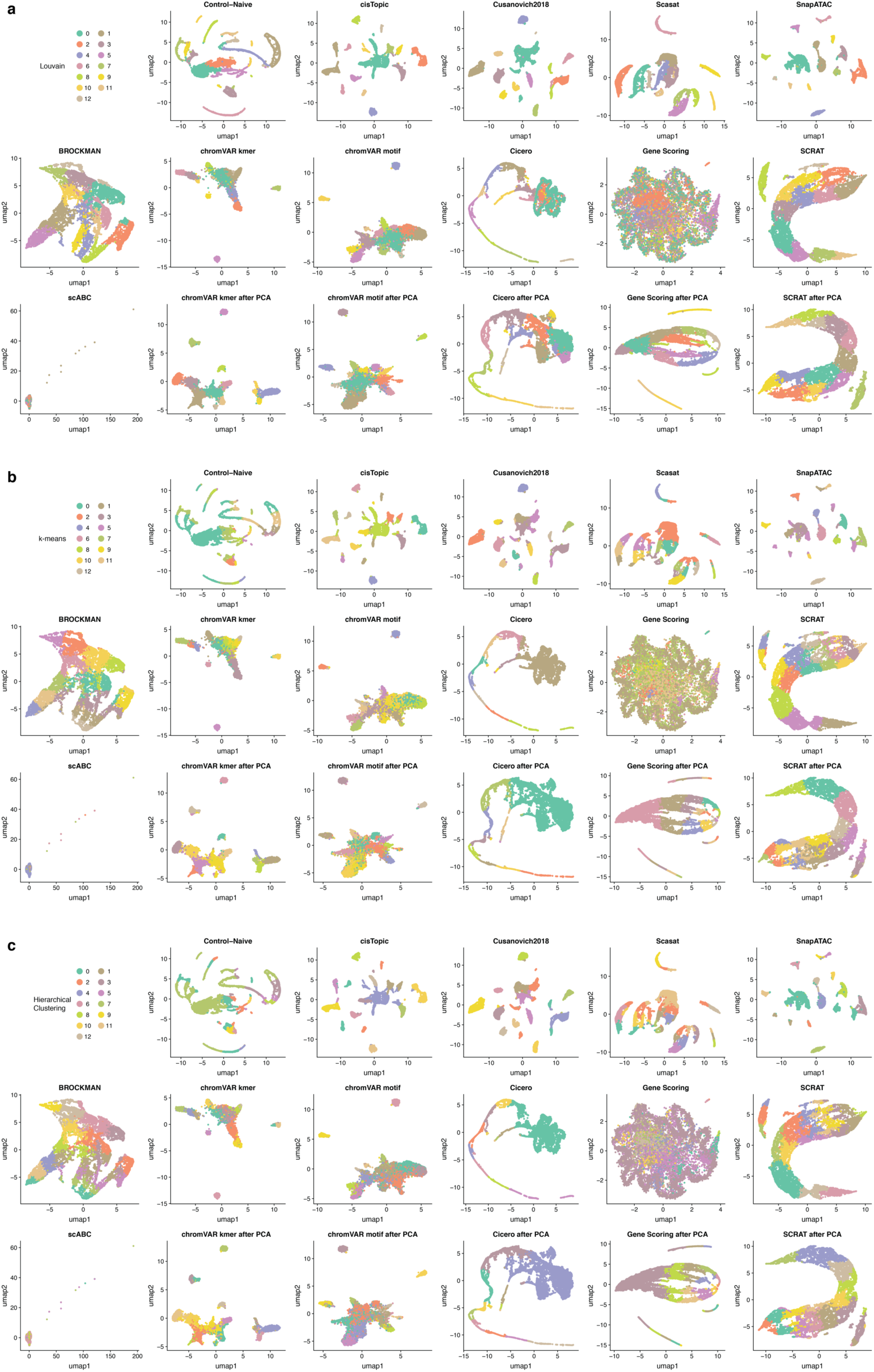
UMAP visualization of cells colored by the clustering solution on the downsampled sci-ATAC-seq mouse dataset using **(a)** the Louvain algorithm, **(b)** k-means clustering, and **(c)** hierarchical clustering (HC).

**Figure S13.**
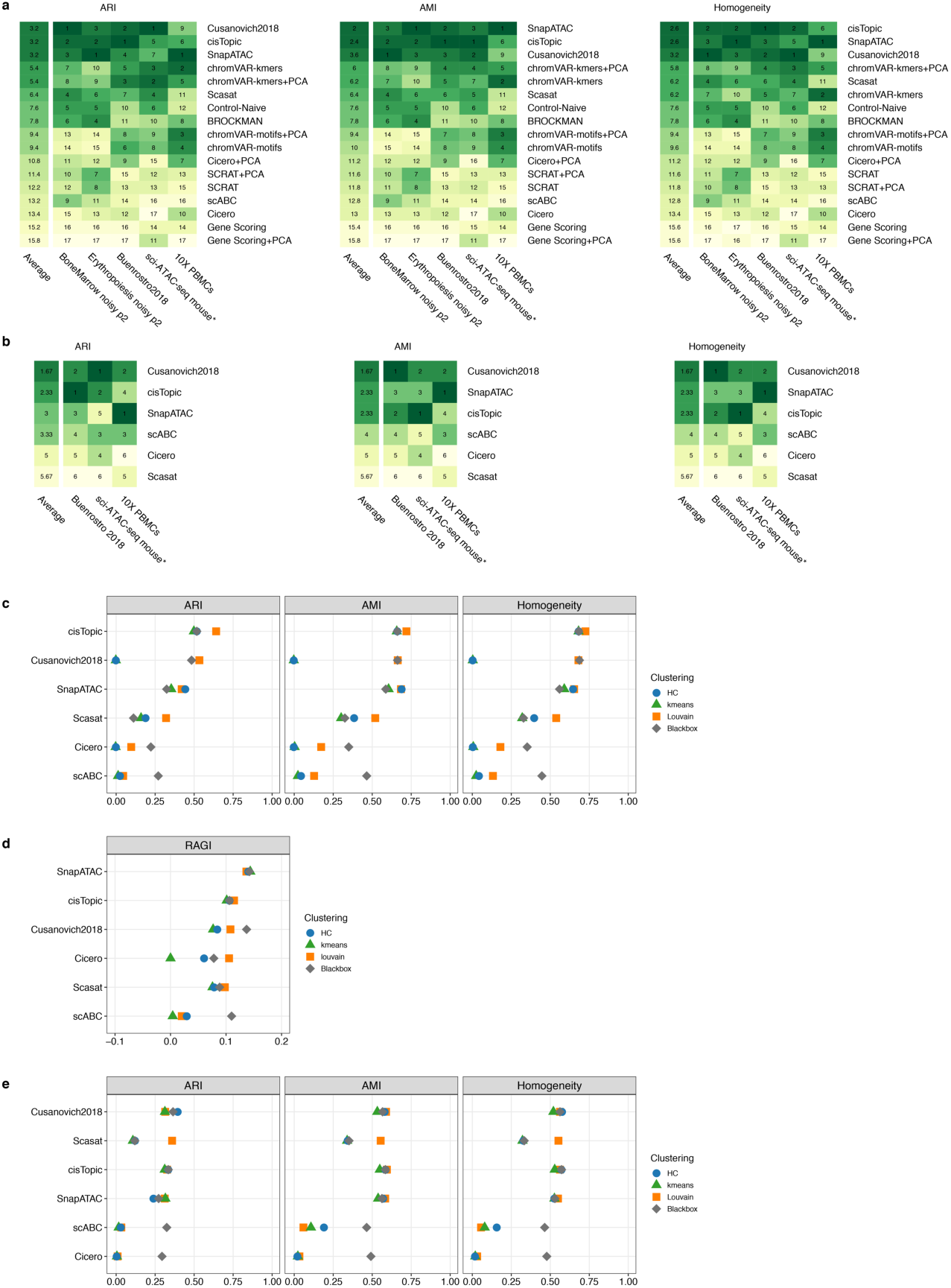
Ranking of method performance. **(a)** Rank was based on the best-performing clustering method for each metric on all methods and datasets. The column on the left shows the averaged rank per method across all datasets. * indicates a downsampled dataset of the indicated original dataset. **(b)** Rank of each method based on the best-performing clustering approach for each metric on methods assessed with an end-to-end clustering pipeline (termed as a ‘blackbox’) applied to the *Buenrostro2018*, downsampled sci-ATAC-seq mouse and 10X PBMCs datasets. The column on the left shows the averaged rank per method over these three datasets. * indicates a downsampled dataset of the indicated original dataset. **(c)** Dot plot of clustering scores for each metric applied to the *Buenrostro2018* dataset, including the ‘blackbox’ approach. **(d)** Dot plot of clustering scores for each metric applied to the 10X PBMCs dataset, including the ‘blackbox’ approach. **(e)** Dot plot of scores for each metric applied to the downsampled sci-ATAC-seq mouse dataset, including the ‘blackbox’ approach.

**Figure S14.**
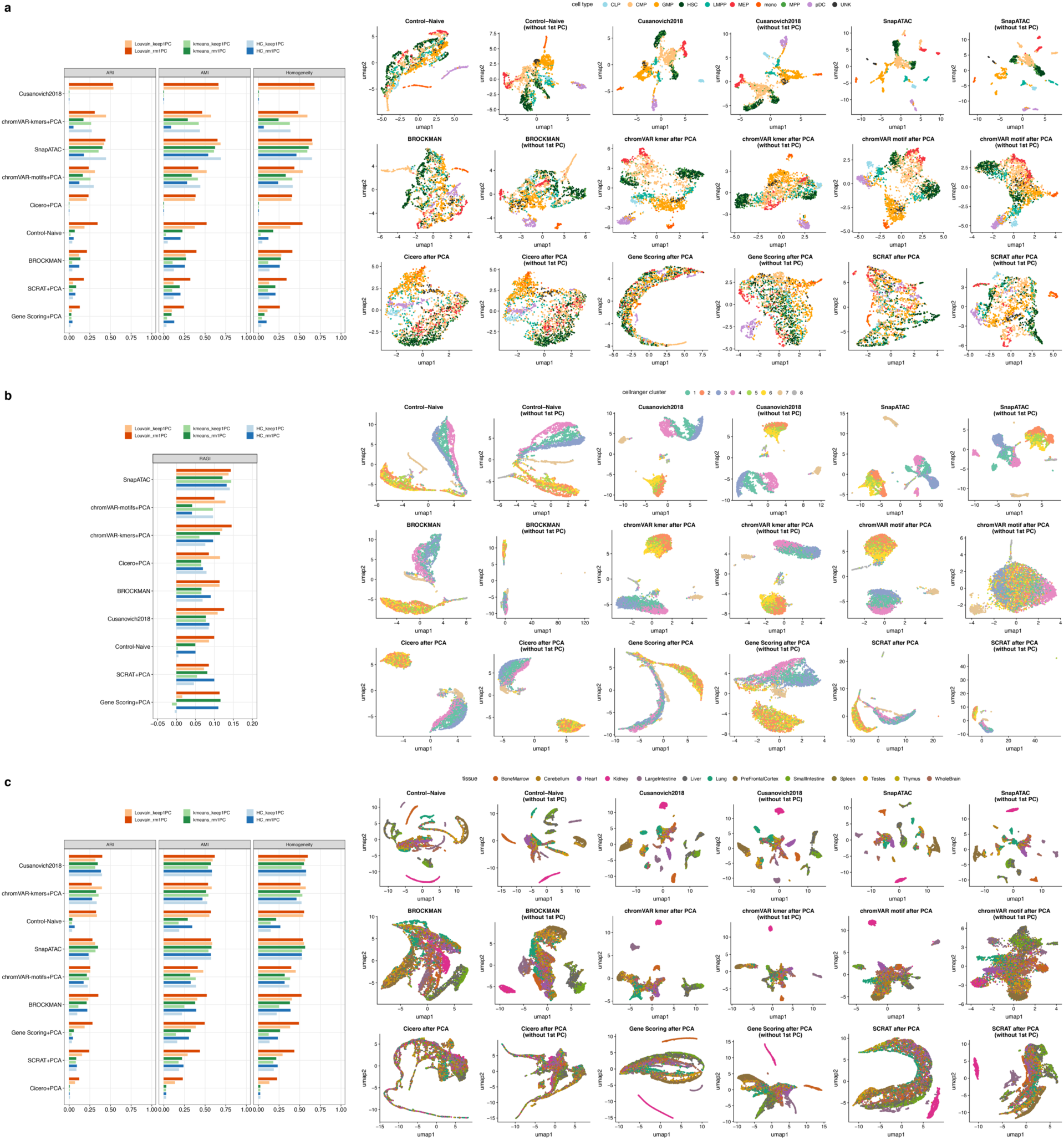
Comparison between keeping the first PC and removing the first PC. ***Left***: Clustering scores when the first PC is kept and for removal of the first PC, for each metric. ***Right:*** UMAP visualization of cells colored by known cell labels. The analyses are performed on **(a)** the *Buenrostro2018* dataset. **(b)** the 10X PBMCs dataset. **(c)** the downsampled sci-ATAC-seq mouse dataset.

**Figure S15.**
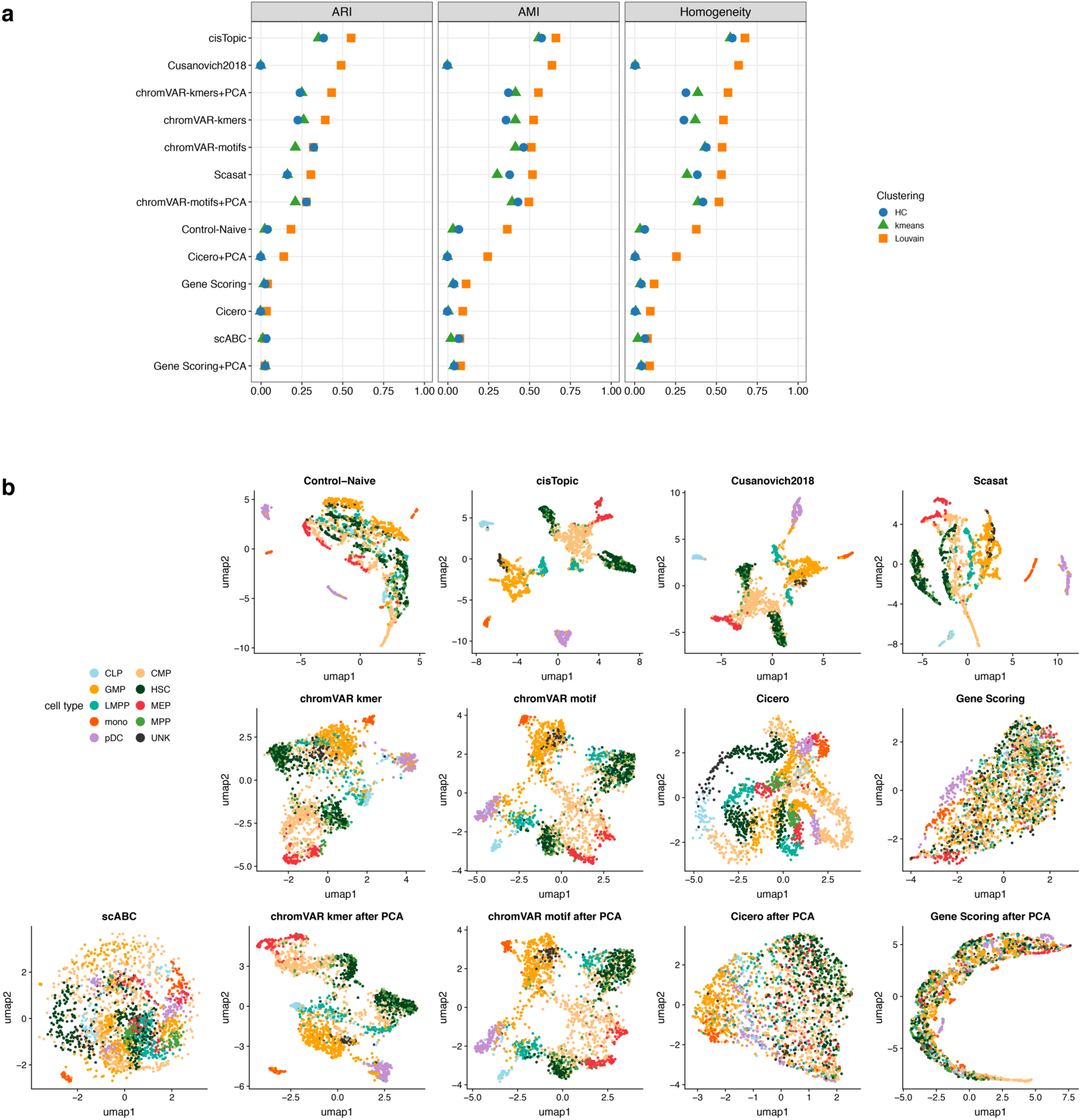
Assessment of methods using the peaks called from bulk ATAC-seq on the *Buenrostro2018* dataset. Only the methods that rely on peaks are included. **(a)** Clustering evaluation according to AMI, ARI and Homogeneity metrics **(b)** UMAP visualization of cells colored by known cell labels.

**Figure S16.**
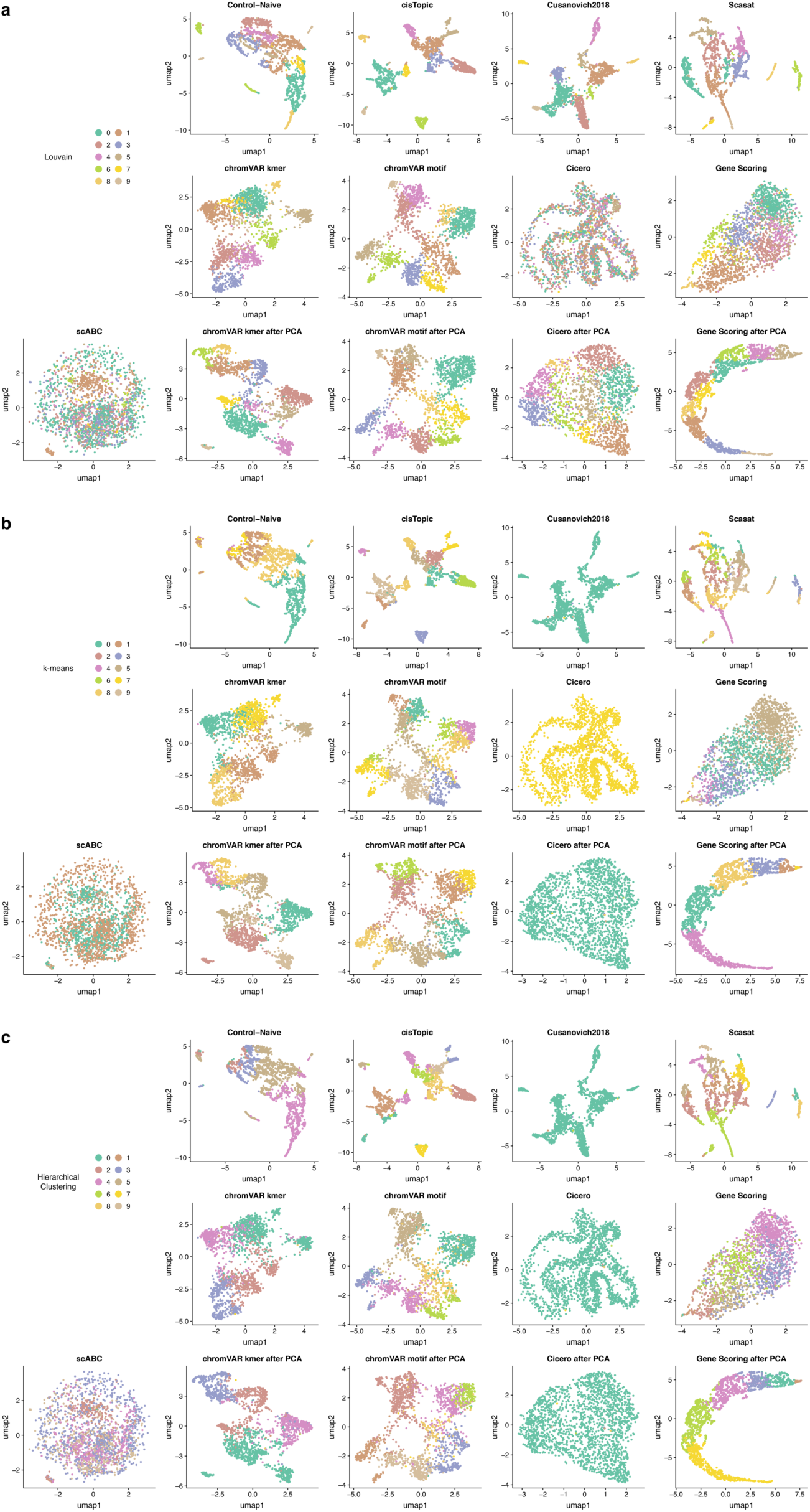
UMAP visualization of cells colored by the clustering solution on the *Buenrostro2018* dataset paired with bulk peaks using **(a)** k-means clustering and **(b)** hierarchical clustering (HC).

**Figure S17.**
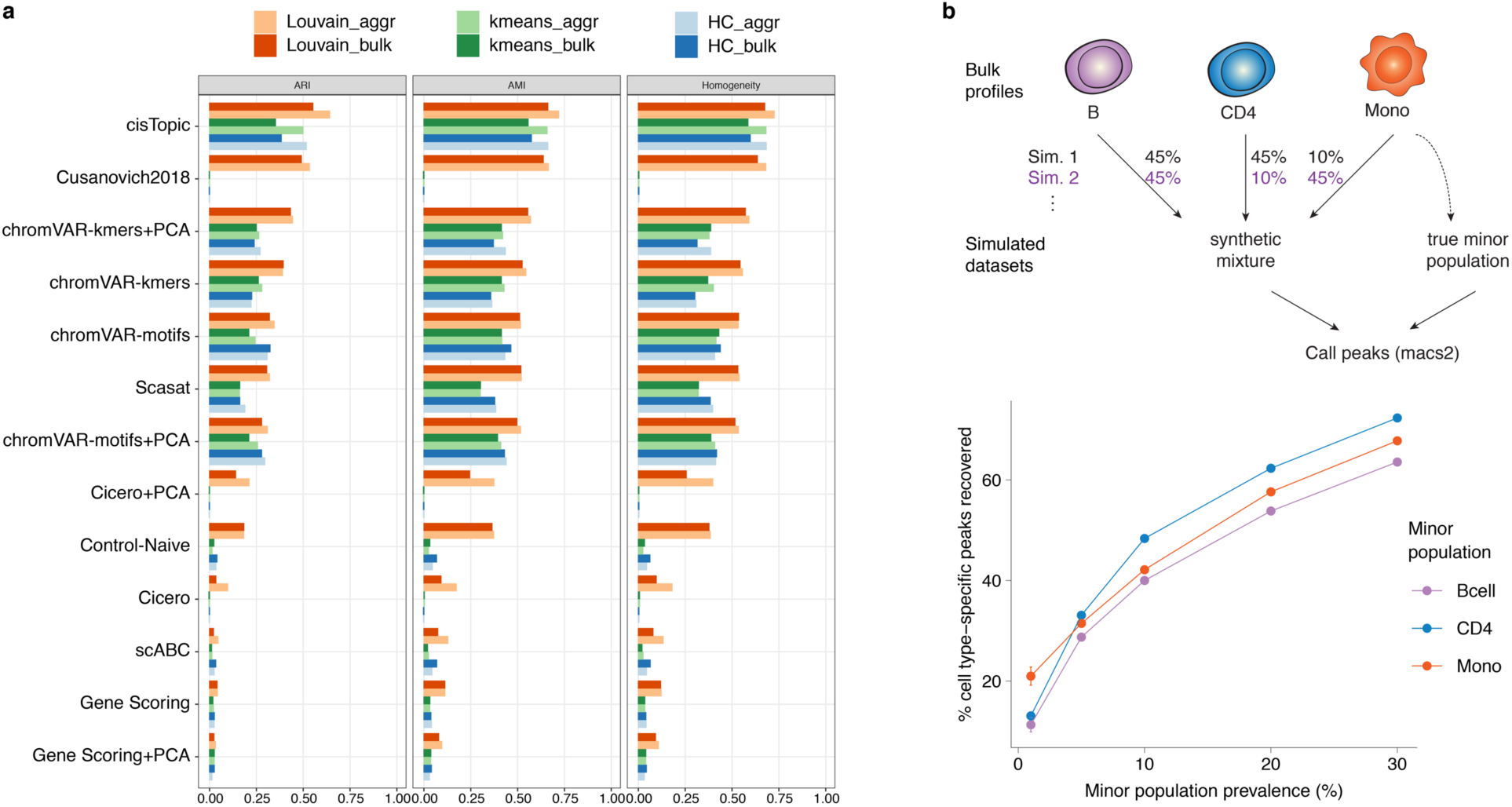
**(a)** Comparison of clustering scores between bulk ATAC-seq peaks and aggregated scATAC-seq peaks for each metric on the *Buenrostro2018* dataset. **(b) Top:** Simulation procedure from bulk ATAC-seq data. The three cell types (B-cells, CD4+ T-cells, and monocytes) are mixed in various proportions for each synthetic mixture. **Bottom:** The results of simulation in (b) **Top**: The x-axis reflects the proportion of the minor population. The y-axis reflects the percentage of recovered cell-type-specific peaks after performing peak calling on each mixture of single cells.

**Figure S18.**
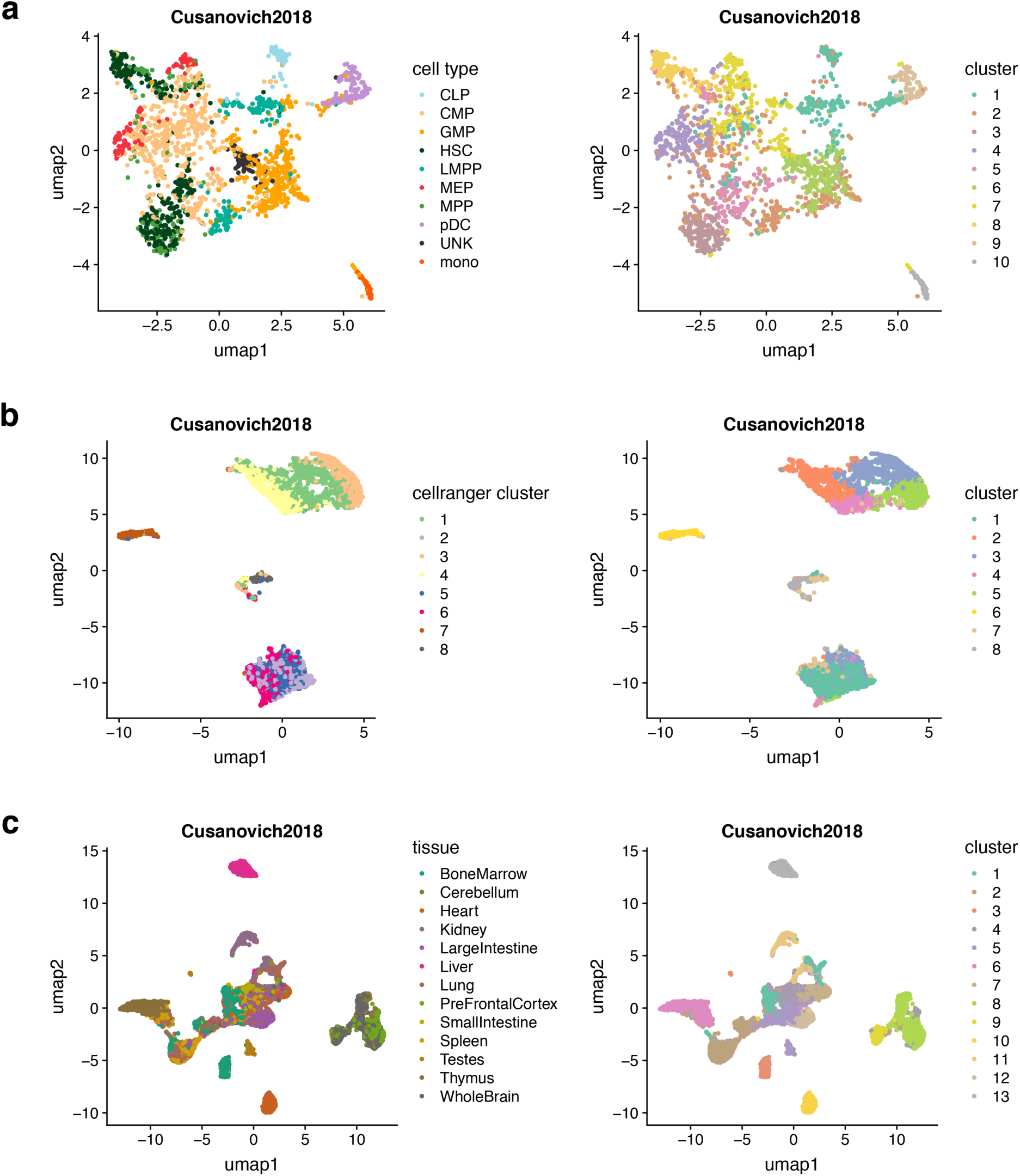
Comparison between the known populations and the identified clades (pseudo-bulk) using *Cusanovich2018*. ***Left***: UMAP visualization of cells colored by the known labels. ***Right:*** UMAP visualization of cells colored by the identified clades using *Cusanovich2018*. The analyses are performed on **(a)** the *Buenrostro2018* dataset. **(b)** the 10X PBMCs dataset. **(c)** the downsampled sci-ATAC-seq mouse dataset.

**Figure S19.**
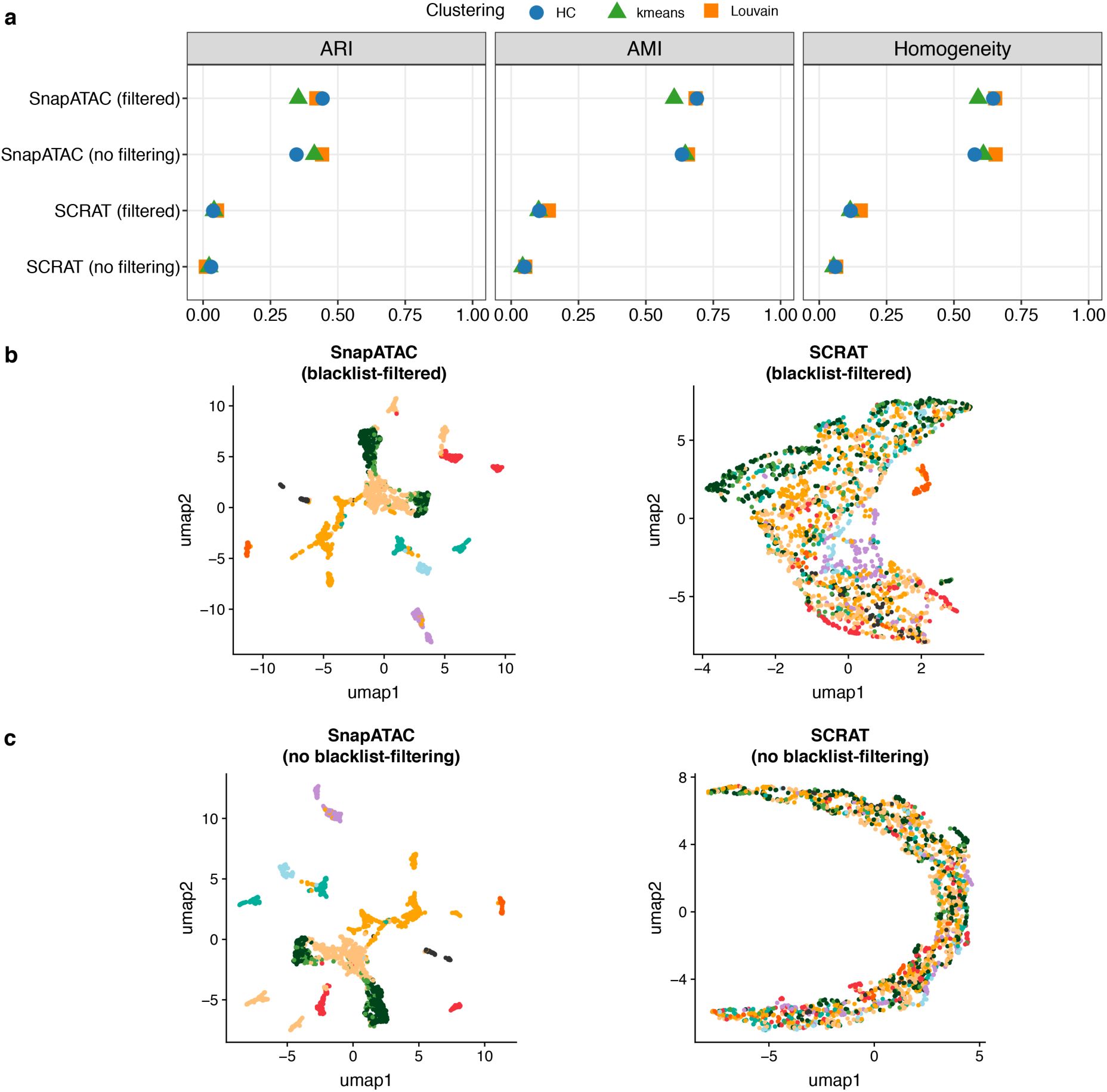
Assessment of the effect of ENCODE blacklisted regions on the benchmarking results in the *Buenrostro2018* dataset. **(a)** Comparison of clustering scores between filtering or not filtering the blacklisted regions **(b)** UMAP visualization based on SnapATAC (***left***) and SCRAT (***right***) feature matrices after filtering the ENCODE blacklisted regions. Cell are colored by the FACS-sorting labels. **(c)** UMAP visualization based on SnapATAC (***left***) and SCRAT (***right***) feature matrices without filtering the ENCODE blacklisted regions. Cell are colored by the FACS-sorting labels.

**Figure S20.**
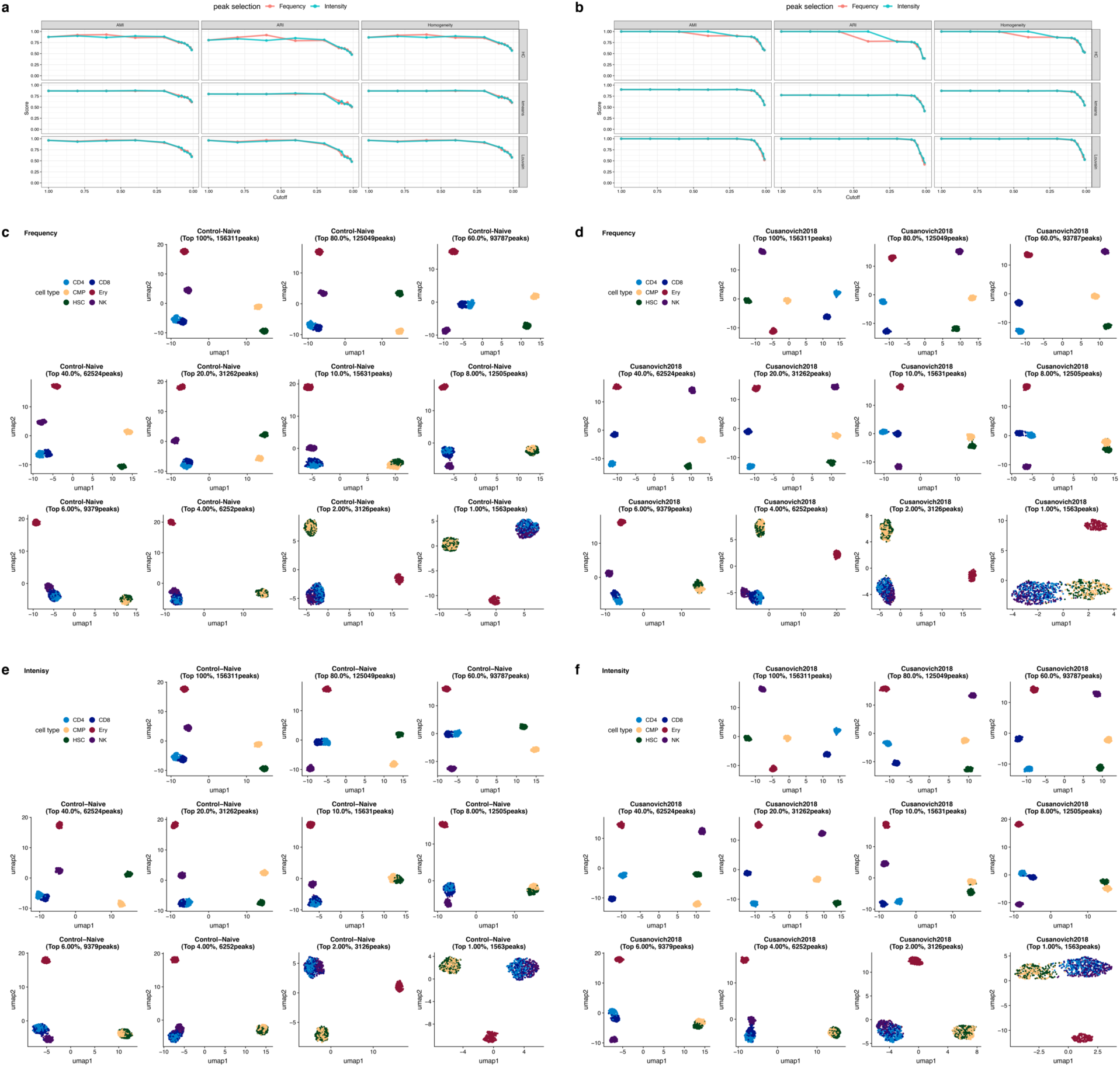
Comparison between frequency-based and intensity-based peak selection for each metric on the simulated bone marrow dataset with a noise level of 0.2 with a coverage of 2,500 fragments. **(a)** Clustering scores for each metric and clustering method across different cutoffs for the Control-Naïve method. **(b)** Clustering scores for each metric and clustering method across different cutoffs for the *Cusanovich2018* method. **(c)** UMAP visualization of cells colored by the known labels using a frequency-based peak selection for Control-naïve method. **(d)** UMAP visualization of cells colored by the known labels using a frequency-based peak selection for *the Cusanovich2018* method. **(e)** UMAP visualization of cells colored by the known labels using an intensity-based peak selection for Control-naïve method. **(f)** UMAP visualization of cells colored by the known labels using an intensity-based peak selection for the *Cusanovich2018* method.

**Figure S21.**
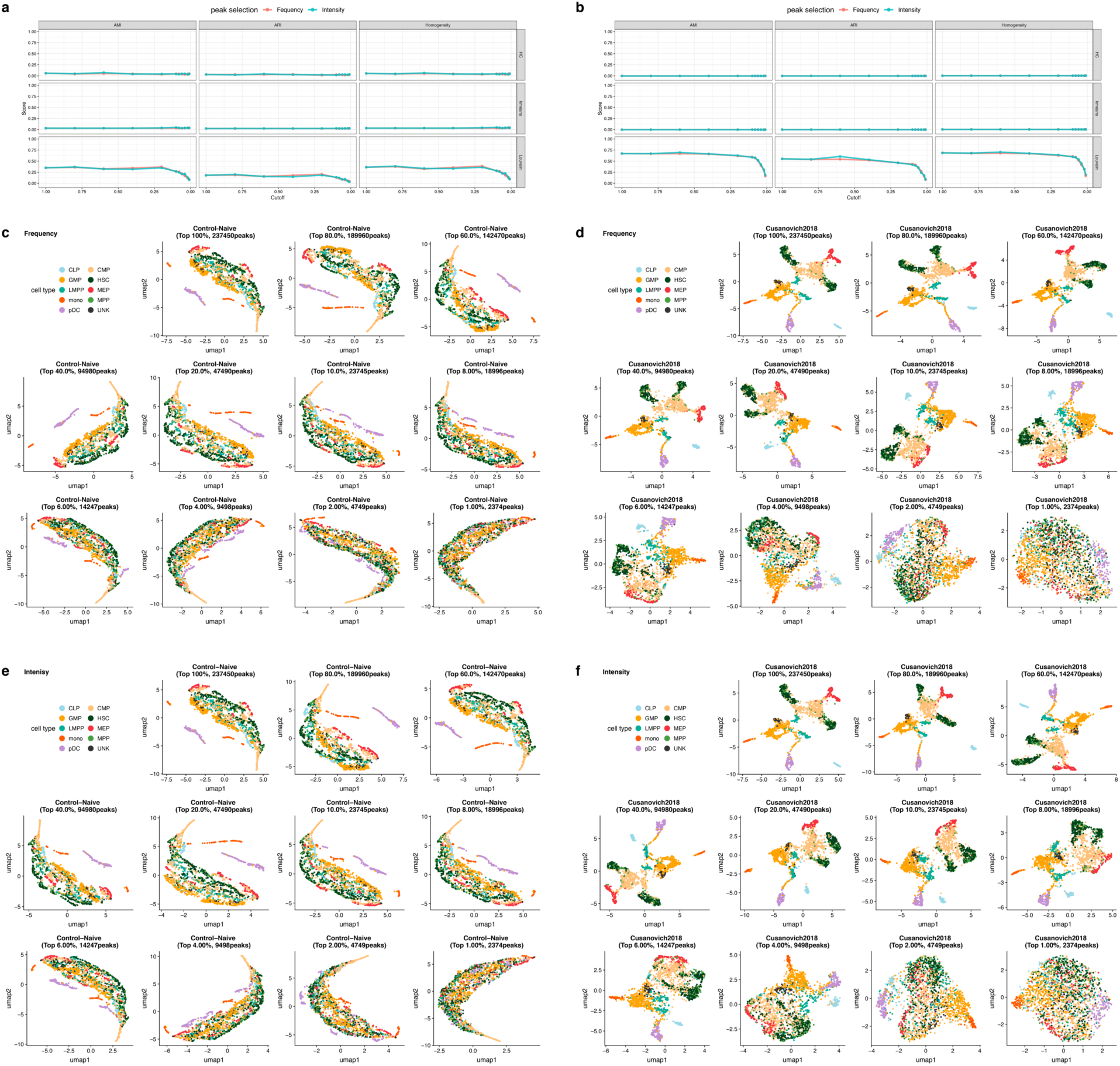
Comparison between frequency-based and intensity-based peak selection for each metric on the *Buenrostro2018* dataset. **(a)** Clustering scores for each metric and clustering method across different cutoffs for the Control-Naïve method. **(b)** Clustering scores for each metric and clustering method across different cutoffs for the *Cusanovich2018* method. **(c)** UMAP visualization of cells colored by FACS-sorting labels using a frequency-based peak selection for the Control-naïve method. **(d)** UMAP visualization of cells colored by FACS-sorting labels using a frequency-based peak selection for the *Cusanovich2018* method. **(e)** UMAP visualization of cells colored by FACS-sorting labels using an intensity-based peak selection for the Control-naïve method. **(f)** UMAP visualization of cells colored by the FACS-sorting labels using an intensity-based peak selection for the *Cusanovich2018* method.

**Figure S22.**
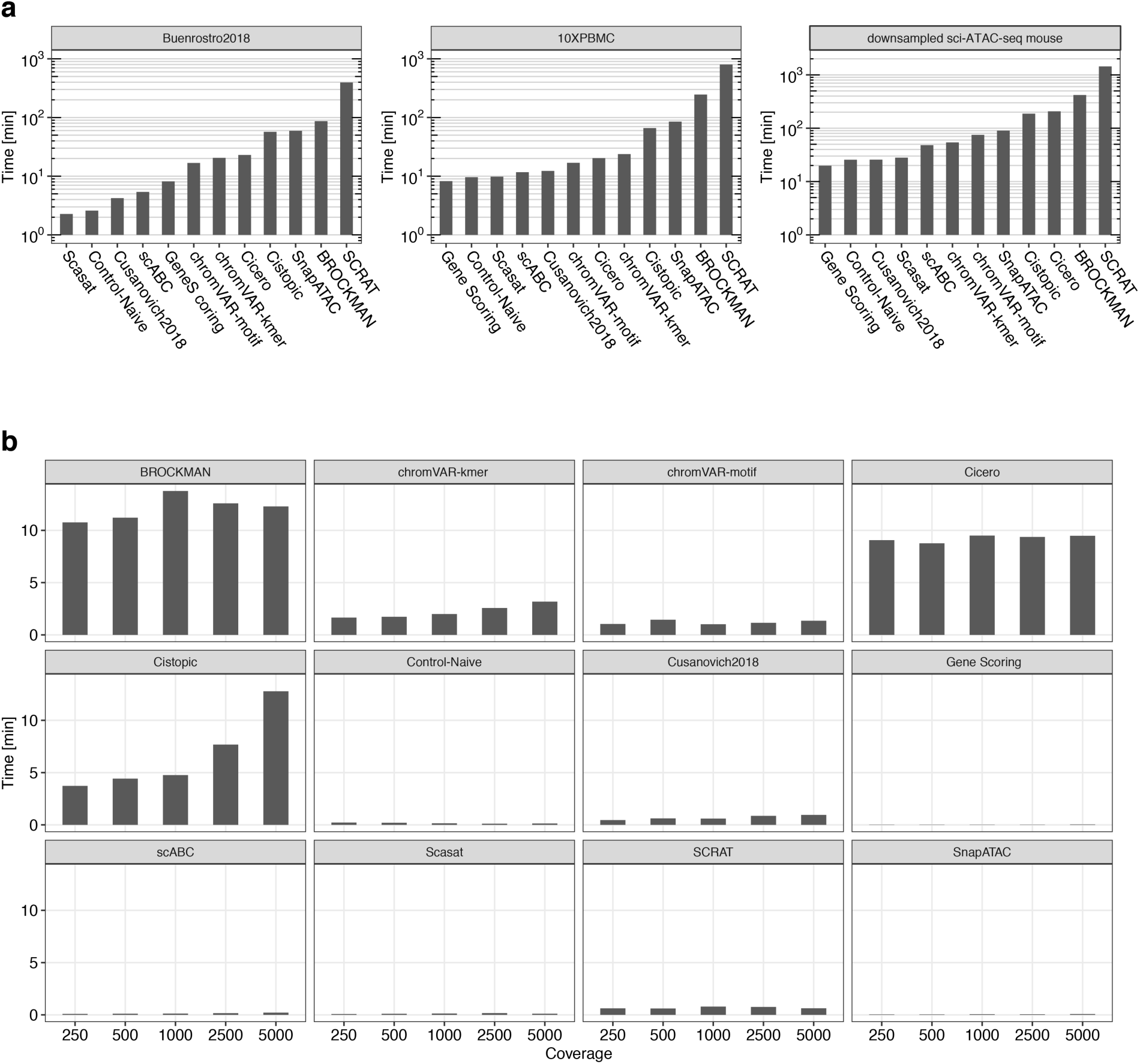
Running time results. **(a)** Running time, in minutes for each method applied to the *Buenrostro2018*, 10X PBMCs, and downsampled sci-ATAC-seq mouse datasets. **(b)** Running time, in minutes for each method on the simulated bone marrow dataset at a noise level of 0.2 with read coverages of 250, 500, 1000, 2500, and 5000 fragments.

